# *C. elegans* E3 ubiquitin ligase EBAX-1 promotes non-apoptotic linker cell-type death through target-directed miRNA degradation

**DOI:** 10.64898/2026.01.07.698237

**Authors:** Lauren B. Horowitz, Olya Yarychkivska, Yun Lu, Shai Shaham

## Abstract

Programmed cell death is essential for animal development and homeostasis, and its disruption accompanies many human disorders. Linker cell-type death (LCD) is a morphologically conserved non-apoptotic developmental cell death program with features resembling polyglutamine-dependent neurodegeneration. In *C. elegans*, LCD execution is mediated by ubiquitin proteasome system (UPS) components, but their proteolytic targets are unknown. Here we demonstrate that EBAX-1/ZSWIM8, a conserved E3 ligase, promotes *C. elegans* LCD by target directed miRNA degradation (TDMD). We show that EBAX-1 acts cell-autonomously as part of the UPS and requires its Cullin-2 binding motif to promote LCD. Loss of *mir-35* family miRNAs, argonautes, or miRNA biogenesis factors, restores LCD to *ebax-1* mutants. Furthermore, expression of *viln-1*/villin mRNA, a predicted *mir-35* target, is upregulated in dying cells and is required for LCD. Together, our studies suggest that TDMD mediated by EBAX-1 is important for the fidelity of non-apoptotic developmental cell death.

## INTRODUCTION

Programmed cell death is a ubiquitous and irreversible metazoan cell fate. In development, cell elimination programs control morphogenesis and tissue remodeling^1^. In mature animals, cell elimination ensures cell number homeostasis and removal of defective cells that can harm the organism, such as autoreactive immune cells or cancerous cells^2^. Aberrant cell death is associated with numerous human disorders, including neurodegenerative diseases, organ infarctions, and cancer^3^. Understanding the molecular mechanisms that direct cell death during development could provide valuable insight into how this process is dysregulated in disease.

Apoptosis is a caspase-dependent cell death mechanism characterized by stereotypical morphological changes, including chromatin condensation and cytoplasm compaction^4–6^. Although apoptosis is observed in many developing animals, and is essential for the development of some, such as the fruit fly *Drosophila melanogaster*^7^, it cannot account for many cell elimination events in mammals^1^. For example, loss of the apoptotic effectors *Apaf1*^8^ or *caspase-9*^9^, or wholesale inhibition of apoptosis in *Bok^-/-^ Bax^-/-^Bak^-/-^* triple mutants^10^, is not sufficient to block murine development. Indeed, these mice can survive to adulthood and exhibit only minor developmental defects^8–10^. Furthermore, although cell death prevention is inherent to the process of malignancy^11^, mutations in core apoptotic genes are not associated with most cancer types^12,13^. Likewise, inhibiting apoptosis effector genes does little to prevent cell loss in mouse models of neurodegeneration^12,13^. Thus, non-apoptotic cell death mechanisms are likely to play major roles in mammalian development and in disease.

Studies of the nematode *C. elegans* uncovered a developmental cell death program, linker cell-type death (LCD), that is morphologically distinct from apoptosis^14^. The *C. elegans* linker cell guides elongation of the male gonad and subsequently dies by LCD to facilitate fusion of the gonadal and cloacal tubes, enabling formation of the sperm exit-channel (**Figure 1a**)^15^. Unlike apoptotic cells, the linker cell displays distinct nuclear envelope crenellations (invaginations), and cytoplasmic organelle swelling at the onset of death^14^. Chromatin condensation is not evident during linker cell death; instead, loss of peripheral heterochromatin is a reproducible feature^14^. As the cell loses contact with its neighbors and is engulfed, it splits asymmetrically, with the larger fragment retaining the nucleus (**Figure 1a**)^16^. While the small fragment is quickly degraded, the larger fragment becomes round and refractile and is degraded through a non-canonical program mediated by RAB-35-dependent removal of ARF-6 from phagocytic membranes^16^. Importantly, linker cell death proceeds independently of all four *C. elegans* caspase-related genes and other apoptotic effectors, as well as regulators of necrosis or autophagy^14,17^.

**Figure 1.**
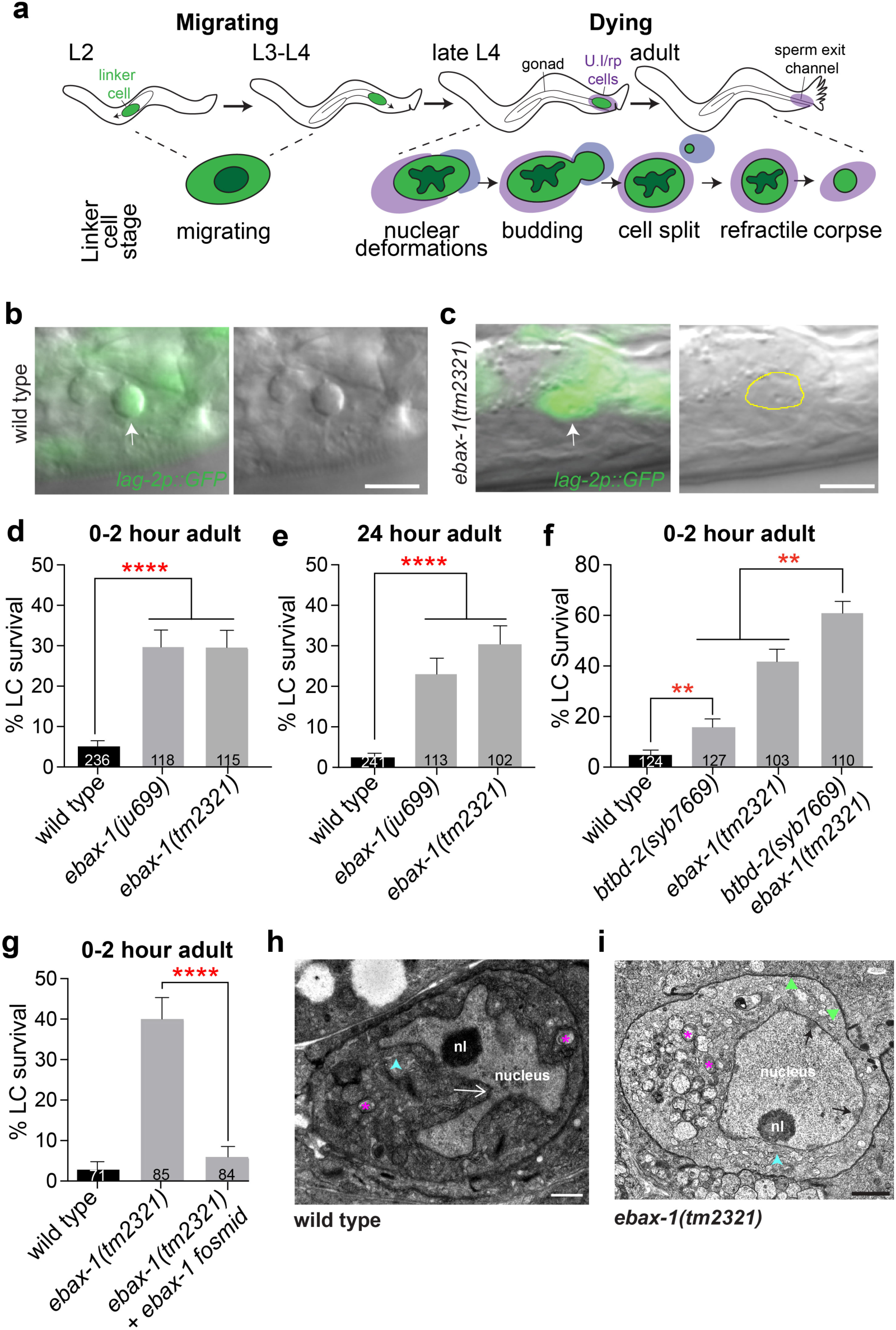
EBAX-1 is required for Linker Cell Death. **(a)** Linker cell migration and morphological changes during death. Adapted from Horowitz and Shaham, 2024^18^. **(b)** Overlaid differential interference contrast (DIC) and fluorescence micrograph (left) of a dying linker cell (arrow) in a wild-type young adult male expressing the linker cell marker *lag-2p::GFP*. The same cell is shown by DIC alone (right) to highlight the refractile morphology of the dying cell. Scale bar, 5 μm. **(c)** Same as **(b)** except showing a surviving linker cell in a young adult *ebax-1(tm2321)* mutant male. Scale bar, 5 μm. **(d and e)** Percent young adult (0-2 hours post L4 molt, d) and 1-day old adults (24 hours post L4 molt, e) with surviving linker cells in mutant males of the indicated genotypes. (****) *p* < 0.0001, Fisher’s exact test. **(f)** Linker cell survival in males of the indicated genotypes. (**) *p* = 0.0061 between wild-type and *btbd-2(syb7669*); (**) *p* = 0.0062 between *ebax-1(tm2321)* and *btbd-2(syb7669) ebax-1(tm2321)*, *p* < 0.0001 between *btbd-2(syb7669)* and *btbd-2(syb7669) ebax-1(tm2321)*; Fisher’s exact test. **(g)** Linker cell survival in males of the indicated genotypes. (****) *p* < 0.0001. Fisher’s exact test. **(h)** Electron micrograph of a dying linker cell in a wild type animal. White arrow, nuclear crenellation. Asterisks, swollen mitochondria. Arrowhead, swollen endoplasmic reticulum. nl, nucleolus. Scale bar, 0.5 μm. **(i)** Electron micrograph of a surviving linker cell in a young adult *ebax-1(tm2321)* mutant animal. Black arrows, heterochromatin. Triangles, adhesion junctions. Asterisks, swollen mitochondria. Arrowhead, swollen endoplasmic reticulum. nl, nucleolus. Scale bar, 1.0 μm. All strains contain *lag-2p::GFP* and *him-5(e1490)*. Number of animals scored is listed inside box. Bar graph data are plotted as mean ± standard error of the proportion.

A molecular framework governing *C. elegans* LCD has been described^18^. Several upstream signaling modules, including two opposing Wnt signaling pathways^19^, a developmental timing pathway involving the *let-7* miRNA and the Zn-finger transcription factor LIN-29^14^, and a MAPKK pathway^20^, function in parallel to activate linker cell death. These pathways act upstream of the heat-shock responsive transcription factor HSF-1, which non-canonically promotes linker cell death^19^. LCD-specific transcriptional targets of HSF-1 include the E2 ubiquitin-conjugating enzyme *let-70*/Ube2D2, whose expression is induced just prior to linker cell death onset^19^. Linker cell expression of additional ubiquitin proteasome system (UPS) components is similarly induced and required for LCD^19^. However, neither the proteolytic targets of the UPS nor the mechanism by which the UPS triggers linker cell demise are known.

LCD-associated morphological changes are evident in dying cells in many developing mammalian tissues, including the nervous system and the male and female reproductive tracts^21^. Cell death accompanied by nuclear crenellations and cytoplasmic organelle swelling is also associated with polyglutamine (polyQ) neurodegenerative disorders, such as Huntington’s disease^22,23^. This association is particularly intriguing given the role of PQN-41, a polyQ-rich protein, in *C. elegans* LCD^20^. Furthermore, TIR-1, a MAPKK scaffold protein, promotes linker cell death^20^, and its *Drosophila* and mouse homologs (Sarm/SARM1) mediate axon degeneration following injury^24^. Thus, proteins related to *C. elegans* LCD effectors promote degenerative processes in mammals, raising the possibility that the molecular program for LCD may be conserved.

Here, we identify EBAX-1, the conserved substrate-recognition subunit of Cullin 2-based E3 ubiquitin ligases, as a key regulator of LCD in *C. elegans*. EBAX-1 is expressed in the linker cell and acts cell autonomously downstream of HSF-1, and together with LET-70/E2-enzyme, to mediate LCD. We show that loss of EBAX-1 promotes inappropriate linker cell survival that can be suppressed by concomitant loss of the argonaute proteins ALG-1 or ALG-2, by linker cell specific loss of miRNA processing enzymes, or by specific depletion of *mir-35* family miRNAs. We further find that loss of a predicted *mir-35* family target mRNA, *viln-1*, encoding a villin-like cytoskeleton protein, results in aberrant linker cell survival and that *viln-1* expression is induced in the linker cell during cell death. Our studies are consistent with previous reports implicating EBAX-1, and its mammalian homolog ZSWIM8, in target-directed miRNA degradation (TDMD), a conserved process regulating miRNA expression^25–27^, and suggest a previously unrecognized role for TDMD in the control of cell death during animal development.

## RESULTS

### *C. elegans* EBAX-1 promotes linker cell death

We previously reported that adult males of a highly mutagenized and fully sequenced *C. elegans* strain, VC40125, generated by the million-mutation project^28^, display inappropriate linker cell survival^19^. Mapping studies suggested that a G-to-T(S82stop) mutation in the *btbd-2* gene, encoding a Cullin-E3 ubiquitin ligase component, may be the causal lesion. Consistent with this, we previously showed that linker-cell-specific expression of a *btbd-2* cDNA restores linker cell death to this strain; however, rescue is incomplete^19^. Thus, another linked mutation may also be involved. Indeed, we noted in VC40125 genomic sequences^28^ a C-to-T (Q1389stop) mutation in the *btbd-2*-linked gene *ebax-1*, encoding a Cullin-2 E3 ubiquitin ligase substrate recognition protein^29^. To determine whether loss of *ebax-1* inhibits linker cell death, we examined two different single mutant strains, *ebax-1(tm2321)* and *ebax-1(ju699)*, containing 551 bp and 1548 bp deletions in the gene, respectively (**Figure 2a**)^29^. Linker cells in wild-type young-adult males are either gone or exhibit a characteristic rounded morphology (**Figure 1b,d**), suggesting loss of adhesion to adjacent tissues and ongoing cell dismantling. By contrast, similarly staged *ebax-1* mutant males often exhibit surviving linker cells that maintain their shape (**Figure 1c,d**), suggesting that cell death fails to initiate in the absence of *ebax-1*. Consistent with this idea, aberrantly surviving linker cells are evident at the same frequency 24 hours after the L4-to-adult transition in both *ebax-1* mutant strains (**Figure 1e**), even though 97% of linker cells in wild-type animals are fully degraded or dying at this stage (n=241). *ebax-1(tm2321) btbd-2(syb7669)* double mutants, carrying a newly generated CRISPR/Cas9 deletion of the *btbd-2* locus, exhibit increased linker cell survival over each single mutant alone (**Figure 1f**), recapitulating the VC40125 strain phenotype^19^. Importantly, a wild-type *ebax-1* genomic transgene fully restores linker cell death to *ebax-1(tm2321)* mutants (**Figure 1g)**, confirming that the *ebax-1(tm2321)* lesion causes linker cell survival. Taken together, these studies reveal that EBAX-1 promotes linker cell death initiation.

**Figure 2.**
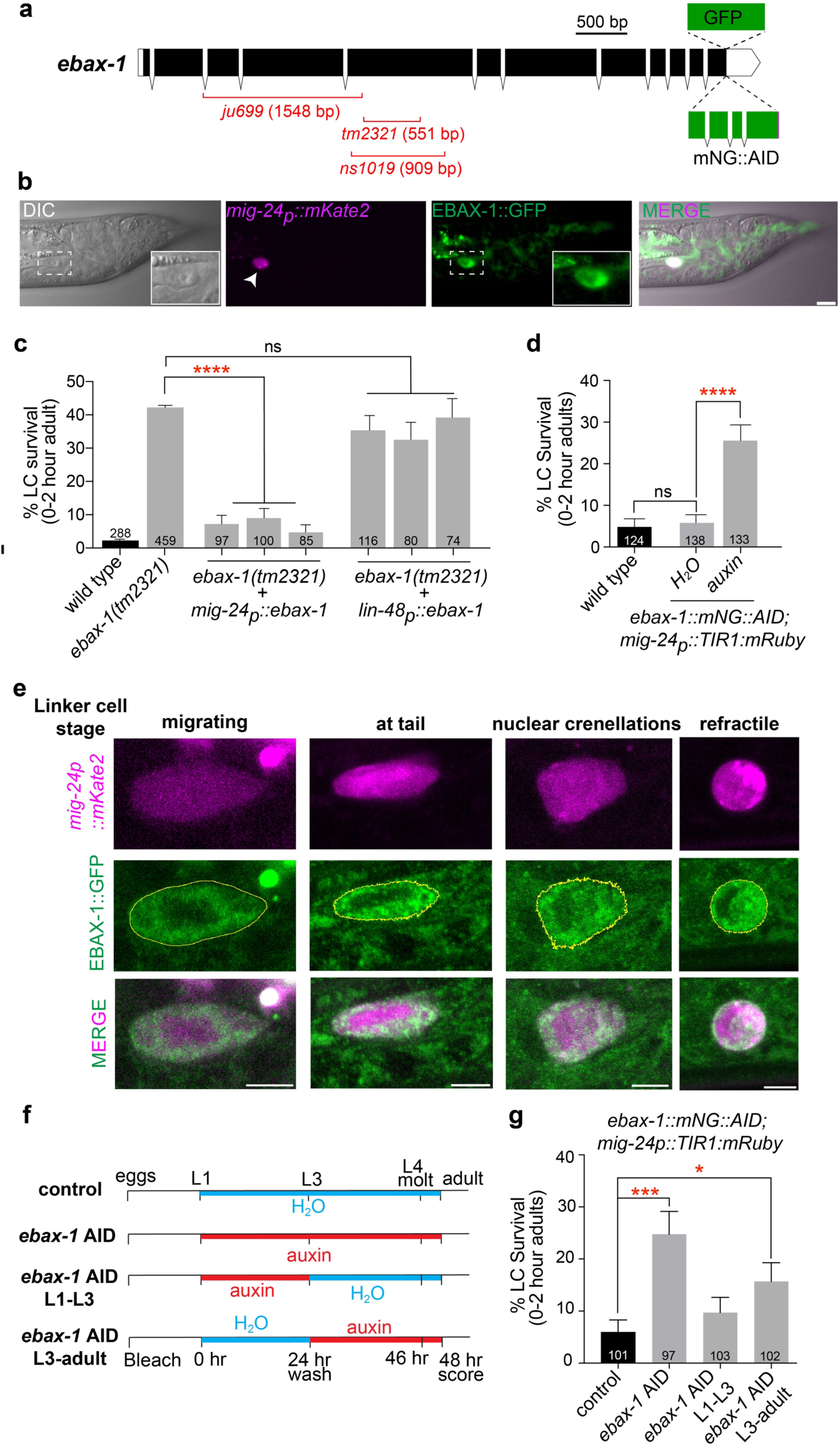
EBAX-1 promotes linker cell death cell autonomously during late larval stages. **(a)** Schematic of the *ebax-1* genomic locus showing mutant alleles and endogenous reporters used in this study. Black boxes, exons. White boxes, UTRs. **(b)** Representative DIC and fluorescence micrographs of an L4-male expressing EBAX-1::GFP (green) and the linker cell marker *mig-24p::mKate2* (magenta). The DIC channel is a single slice; fluorescence channels are maximum-intensity (mKate2) and sum-intensity (GFP) projections of a background-subtracted z-stack. Insets, EBAX-1::GFP localization in the linker cell. Scale bar, 5 μm. **(c)** Linker cell survival in indicated genotypes. *mig-24* and *lin-48* promoters are expressed in the linker cell and U.I/rp engulfing cells, respectively. Three independent lines tested for each rescue construct. (****) *p* < 0.0001; ns, not significant, from left to right: *p* = 0.2969, 0.4519, 0.7376; Fisher’s exact test. **(d)** Linker cell survival in wild-type and in animals expressing *ebax-1::mNeonGreen::AID* as well as *mig-24p::TIR1. ebax-1::mNG::AID; mig-24p::TIR1* mutant animals were raised on water or on auxin to deplete EBAX-1 in the linker cell. (****) *p* < 0.0001; ns, not significant, *p* = 0.7892; Fisher’s exact test. **(e)** Representative confocal maximum intensity projections of wild-type linker cells expressing EBAX-1::GFP at indicated linker cell stages (see Figure 1A). Scale bar, 3 μm. **(f)** Timeline of EBAX-1 auxin inducible degradation experiment, showing conditions relevant for **g**. **(g)** Linker cell survival in *ebax-1::mNG::AID; mig-24p::TIR1* mutant animals raised on water or auxin according to the conditions described in (F). (***) *p* = 0.0003; (*) *p* = 0.0402; Fisher’s exact test. Bar graph data are plotted as mean ± standard error of the proportion. Number of animals scored listed inside box.

### EBAX-1 is required for nuclear changes during LCD

Linker cell death is accompanied by characteristic ultrastructural changes, including deep nuclear envelope crenellations, loss of nuclear periphery-associated heterochromatin, and swelling of mitochondria and endoplasmic reticulum (**Figure 1h)**^14,21^. Using serial-section electron microscopy, we found that although surviving linker cells in *ebax-1(tm2321)* mutants show mitochondrial and endoplasmic reticulum changes characteristic of LCD (**Figure 1i, asterisk, arrowhead)**, nuclear invaginations are not observed, heterochromatin is maintained, and adhesion junctions to surrounding cells remain intact (0-2 hours, n=3; 24 hours, n=1; **Figure 1i**). These ultrastructural features resemble those we previously described in *pqn-41* mutants^20^. Thus, *ebax-1* and *pqn-41* both control nuclear, but not cytoplasmic, changes accompanying linker cell death, suggesting that separate genetic pathways drive cell nucleus and organelle dynamics, and that inhibition of the nuclear pathway is sufficient to prevent linker cell death and degradation.

### EBAX-1 functions cell autonomously in late larval stages for linker cell death

To determine where EBAX-1 functions to effect linker cell death, we used CRISPR/Cas9^30^ to insert *gfp* coding sequences immediately upstream of the *ebax-1* genomic stop codon (**Figure 2a**). The resulting EBAX-1::GFP fusion protein does not perturb linker cell death (**Supplementary Fig. 1**), and localizes to the linker cell, among other cells (**Figure 2b**). EBAX-1::GFP fluorescence is predominantly confined to the linker cell cytoplasm and excluded from the nucleus (**Figure 2b, inset**), suggesting that EBAX-1 may function in the linker cell cytoplasm to drive nuclear changes during LCD.

To confirm that EBAX-1 functions in the linker cell, we introduced a *mig-24p::ebax-1* cDNA transgene, which drives expression of *ebax-1* specifically in the linker cell, into *ebax-1(tm2321)* mutants and assessed linker cell survival. While 44% of *ebax-1(tm2321)* mutant males exhibit linker cell survival 0-2 hours post the larva-to-adult molt, only 7% of *ebax-1(tm2321)* mutants carrying the *mig-24p::ebax-1* transgene display surviving linker cells (**Figure 2c**, 3 transgenes scored). Furthermore, a *lin-48p::ebax-1* cDNA transgene, which promotes expression of *ebax-1* in the U.I/rp cells that engulf the linker cell corpse^16^, fails to rescue inappropriate linker cell survival in *ebax-1* mutants (**Figure 2c**). Thus, EBAX-1 functions cell autonomously to promote linker cell death.

EBAX-1::GFP fusion protein is first evident in the linker cell in third larval stage (L3) animals, and the protein remains detectable until the cell is degraded (**Figure 2e**, n=24 animals). To determine if EBAX-1 function is indeed required during this time period, we used CRISPR/Cas9 to insert auxin-inducible degron (AID) sequences^31–33^ into the endogenous *ebax-1* locus (**Figure 2a**). Animals harboring this *ebax-1::mNeonGreen::AID* gene fusion and also expressing a *mig-24p::TIR1* transgene, encoding an *Arabidopsis* E3 ligase protein that promotes auxin-dependent degradation of AID-tagged proteins^31–33^, display aberrant linker cell survival under continual auxin incubation (**Figure 2d**), consistent with EBAX-1’s cell autonomous function. Auxin exposure between the L1 and L3 stages only (**Figure 2f**), does not affect linker cell death (**Figure 2g**). However, auxin exposure after the L3 stage results in significant linker cell survival **(Figure 2g**). We also examined the linker cell in animals expressing TIR1 using an *eft-3* ubiquitous promoter^32^, and observed similar results, although linker cell survival is less pronounced in these experiments (**Supplementary Fig. 2a,b**). Consistent with this reduced penetrance, auxin lowers, but does not eliminate, EBAX-1::mNeonGreen::AID signal in these animals (compare **Supplementary Fig. 2c and 2d**). Taken together, our results suggest that EBAX-1 function after the L3 larval stage is required for linker cell death.

### EBAX-1 functions as a Cullin 2-associated E3 ubiquitin ligase during LCD

EBAX-1 protein contains several conserved domains. These include N-terminal BC-box and Cul2-box motifs, which bind Elongin B/C and Cullin 2 respectively^29,34^; SWIM and A domains, whose functions are less understood^29,35^; and a C-terminal region predicted to mediate substrate binding^29^ (**Figure 3a,b**). To determine which of these domains is required for linker cell death, we introduced *mig-24p::ebax-1* cDNA transgenes encoding EBAX-1 variants lacking individual domains into *ebax-1(tm2321)* mutants and assessed linker cell death restoration (**Figure 3b-d**). Expression of either the N-terminal half (amino acids 1-852) or C-terminal half (amino acids 853-1717) alone fails to rescue *ebax-1(tm2321)* mutants (**Figure 3c**). Likewise, deletion of the Cul2-box, SWIM, or A domains abolishes EBAX-1 LCD activity (**Figure 3d**). By contrast, deletion of the BC-box does not affect EBAX-1 activity (**Figure 3d**). Thus, the CUL-2, SWIM and A domains are required for EBAX-1 function during LCD while the BC-box domain is not.

**Figure 3.**
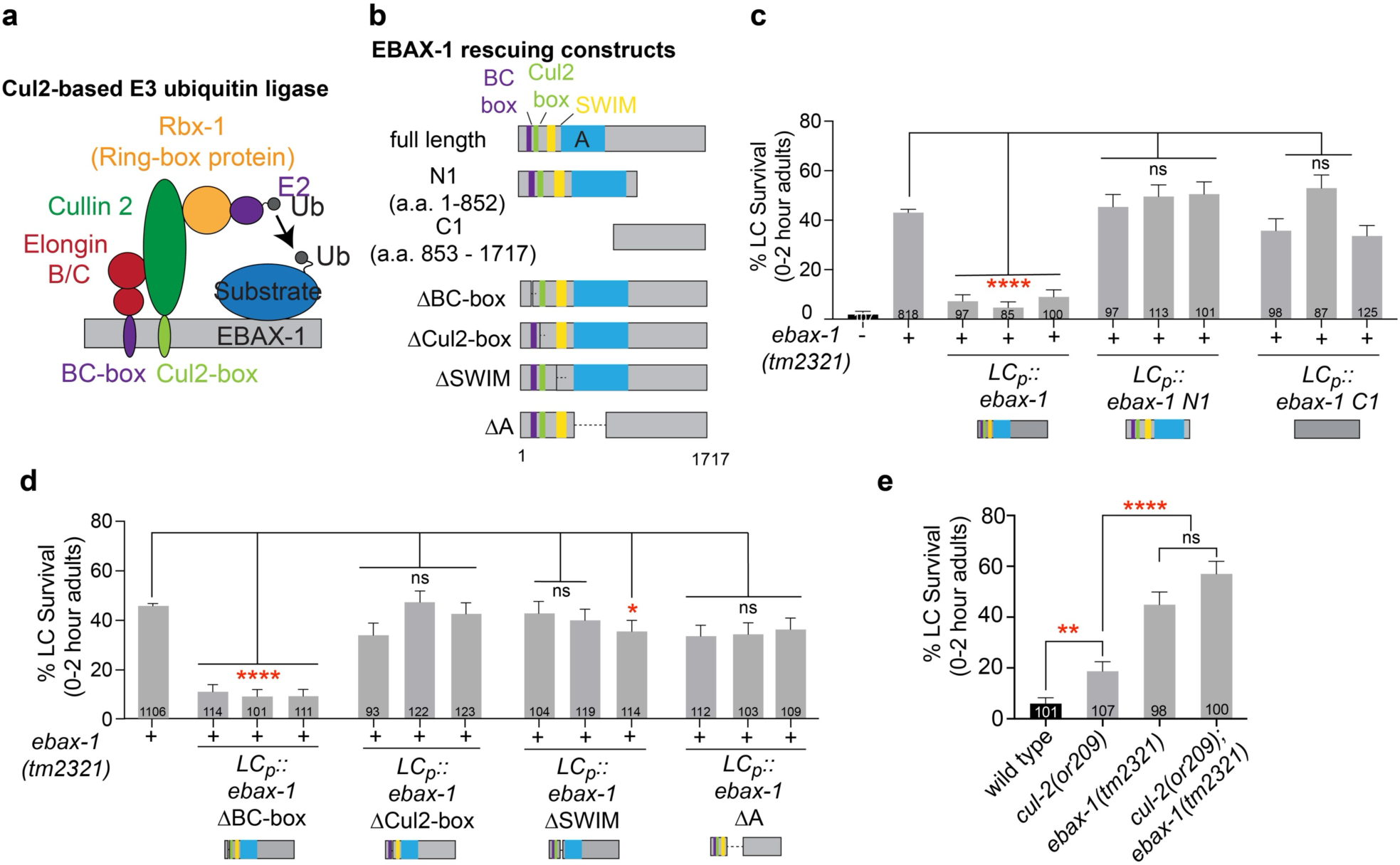
EBAX-1 functions as a substrate recognition subunit of a Cul2-based E3 ligase during linker cell death. **(a)** EBAX-1 functions as a BC-box-type Cullin-2 RING E3 ubiquitin ligase. Adapted from Wang et al., 2013^29^. **(b)** Schematic of EBAX-1 domain structure and deletion constructs used in **(c, d)**. a. a., amino acid. **(c and d)** Linker cell survival in indicated genotypes. Rescuing constructs shown in **(b)** are driven by the linker cell (LC) specific *mig-24* promoter. Three independent lines were tested for each construct. Data shown for the complete linker cell-specific *ebax-1* cDNA rescue, *LCp::ebax-1*, is reprinted from Figure 2C. (****) *p* < 0.0001, (*) *p* = 0.0443, ns, not significant from left to right for *mig-24p::ebax-1 N1*: *p* = 0.8808, 0.6613, 0.6612, for *mig-24p::ebax-1 C1: p* = 0.3267, 0.0787, 0.7801, for *mig-24p::ebax-1 ΔCul2-box*: *p* = 0.2328, 0.8973, 0.3321, for *mig-24p::ebax-1 ΔSWIM*: *p =* 0.6671, 0.4785, for *mig-24p::ebax-1* ΔA: *p* = 0.0701, 0.6539, 0.1094; Fisher’s exact test. **(e)** Linker cell survival in indicated genotypes. (****) *p* < 0.0001; (**) *p* = 0.0061; ns, not significant *p* = 0.1176; Fisher’s exact test. Bar graph data are plotted as mean ± standard error of the proportion. Number of animals scored listed inside box.

To determine whether the requirement for the Cul-2 box reflects a requirement for Cullin 2, we examined linker cell death in weak *cul-2(or209)* loss-of-function mutants and noted a significant increase in linker cell survival (**Figure 3e**). Moreover, inappropriate linker cell survival in *cul-2(or209); ebax-1(tm2321)* double mutants is not significantly different from linker cell survival in *ebax-1* single mutants (**Figure 3e**). These findings indicate that EBAX-1 and CUL-2 function in the same genetic pathway for linker cell death. Furthermore, previous studies showing binding of EBAX-1 to CUL-2 in a different context^29^ are consistent with the idea that binding to CUL-2 likely mediates EBAX-1 killing activity.

### EBAX-1 acts downstream of HSF-1 to promote linker cell death

We previously showed that the E2 ubiquitin conjugating enzyme LET-70, a component of the UPS homologous to mammalian UBE2D2, functions downstream of HSF-1 to promote linker cell death^19^. We wondered, therefore, whether EBAX-1, another UPS component, functions similarly. Although *hsf-1(sy441); ebax-1(tm2321)* double mutants, harboring a partial loss of function allele of *hsf-1*, have enhanced linker cell survival (**Figure 4a**), this finding is not sufficient to determine the relative order of action of these two genes in the LCD pathway. Instead, we assessed linker cell survival in *ebax-1(tm2321)* mutants that also contain a genomically integrated single-copy homozygous transgene expressing HSF-1(R145A), an *hsf-1* gain-of-function allele that promotes linker cell death^19^. We found that whereas inappropriate linker cell survival caused by mutation of the *sek-1* MAPKK gene is alleviated by HSF-1(R145A), indicating that SEK-1 acts upstream of HSF-1^19^, linker cell survival in *ebax-1(tm2321)* mutants is not blocked by this HSF-1 variant (**Figure 4b**). Thus, like LET-70, EBAX-1 likely acts downstream of HSF-1 (**Figure 4d)**. Furthermore, double mutants between *ebax-1(tm2321)* and animals deficient for the Wnt signaling pathway (*bar-1(RNAi*)^19^) or the MAPK pathway (*pqn-41(ns294)*^20^) do not have excess linker cell survival compared to *ebax-1(tm2321)* single mutants alone (**Supplementary Fig. 3**), consistent with the idea that these genes act in the same pathway and that *ebax-1* functions downstream of both (**Figure 4d)**.

**Figure 4.**
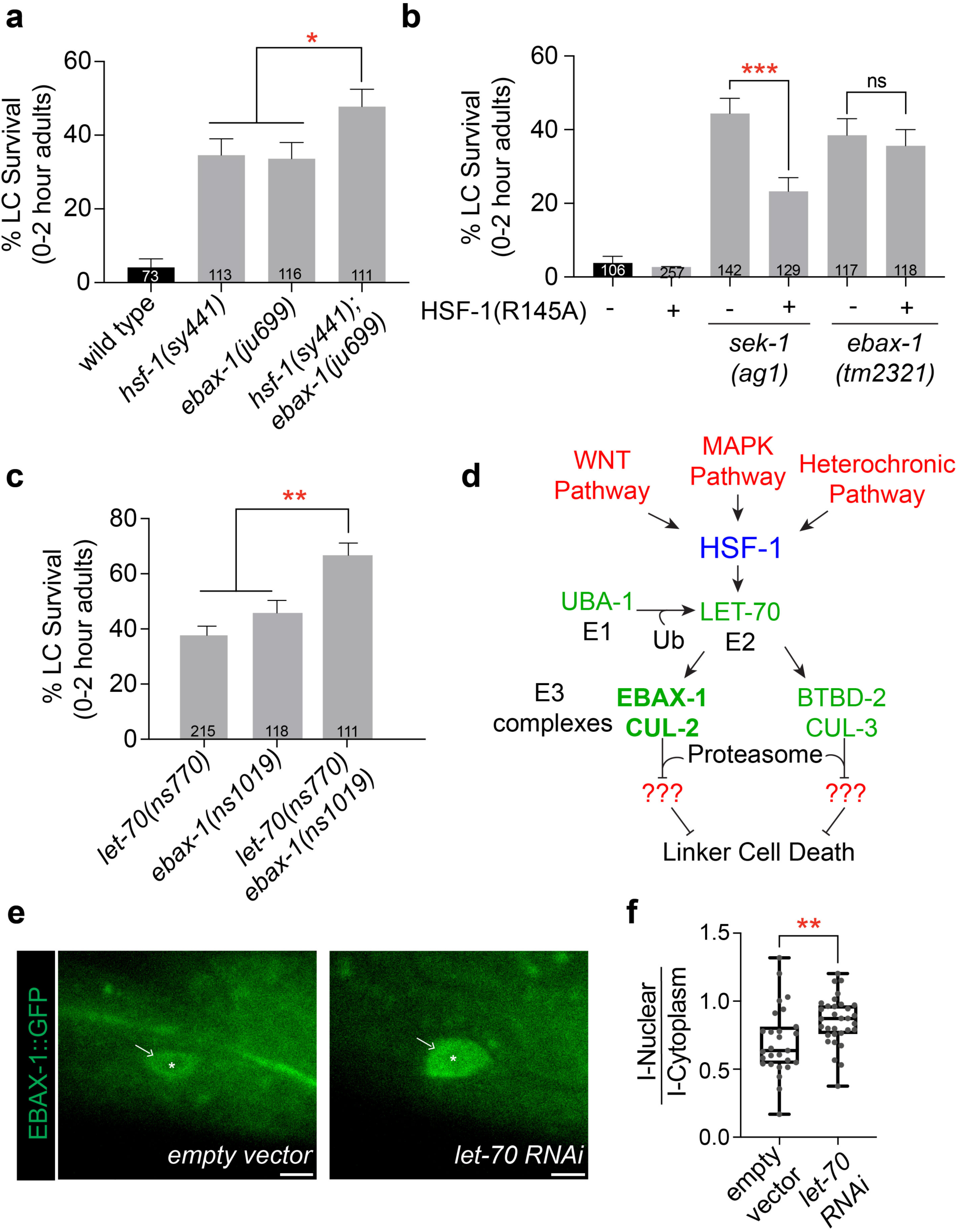
EBAX-1 functions downstream of HSF-1 as part of the ubiquitin proteasome system to effect linker cell death. **(a)** Linker cell survival in indicated genotypes. (*) *p* = 0.0317 between *ebax-1(ju699)* and *hsf-1(sy441); ebax-1(ju699)* and *p* = 0.0428 between *hsf-1(sy441)* and *hsf-1(sy441); ebax-1(ju699)*; Fisher’s exact test. **(b)** Linker cell survival in indicated genotypes. (***) *p* = 0.0003; ns, not significant *p* = 0.6864, Fisher’s exact test. **(c)** Linker cell survival in indicated genotypes. (**) *p* = 0.0021, between *ebax-1(tm2321)* and *let-70(ns770) ebax-1(tm2321)* and *p <* 0.0001 between *let-70(ns770)* and *let-70(ns770) ebax-1(tm2321)*, Fisher’s exact test. **(d)** Proposed placement of EBAX-1 and CUL-2 in the linker cell death signaling pathway. **(e)** Representative fluorescence micrographs of EBAX-1::GFP localization in the linker cell of wild-type animals treated with empty vector or *let-70 RNAi*. Images are sum projections of three consecutive z-slices (∼1.0 μm). Asterisk, nucleus. Arrow, cytoplasm. Scale bar, 5 μm. **(f)** Box and whisker plots of the ratio of EBAX-1::GFP pixel intensity in the linker cell nucleus (I-Nuclear) to the pixel intensity in the cytoplasm (I-Cytoplasm) from wild-type animals treated with empty vector or *let-70* RNAi. Each dot represents one cell. Boxes, interquartile range. Horizontal lines, median. Whiskers, 95% confidence interval. (**) *p* = 0.0083, unpaired t-test, two tailed, t = 2.735, degrees of freedom = 57. Bar graph data are plotted as mean ± standard error of the proportion. Number of animals scored listed inside box.

### LET-70 prevents EBAX-1 nuclear localization

To determine if LET-70 functions exclusively with EBAX-1, or in conjunction with other E3 complexes, we used CRISPR/Cas9 to generate a 909-bp putative null *ebax-1* deletion allele (*ebax-1(ns1019)*; **Figure 2a**) in wild-type animals and in animals containing the tightly linked hypomorphic *let-70(ns770)* mutation, which results in a P61S substitution in the LET-70 protein^19^. *ebax-1(ns1019)* single mutants exhibit linker cell survival comparable to other *ebax-1* alleles (**Figure 4c**). However, *ebax-1(ns1019) let-70(ns770)* double mutants have significantly higher linker cell survival than either single mutant alone (**Figure 4c**). This result is in line with our findings that *ebax-1(tm2321) btbd-2(syb7669)* double mutants have enhanced linker cell survival, and suggests that LET-70 likely acts with E3 proteins besides EBAX-1, such as BTBD-2 (**Figures 1f**, **4d**).

To further test the relationship between EBAX-1 and LET-70, we examined EBAX-1::GFP localization when *let-70* is inactivated. Animals expressing EBAX-1::GFP were treated with *let-70* RNAi, which blocks linker cell death^19^, and imaged at the onset of linker cell death. We found that *let-70* knockdown significantly increases EBAX-1::GFP nuclear localization (**Figure 4e, f**). Thus EBAX-1 localization depends on LET-70 and the two proteins likely function together, in one of multiple E3 complexes, to promote linker cell death (**Figure 4d**).

### EBAX-1 likely promotes linker cell death through target-directed miRNA degradation

EBAX-1 is similar in sequence to mammalian ZSWIM8, which mediates target-directed miRNA degradation (TDMD), a process regulating miRNA stability^25,26,36^ (**Figure 5a**). In TDMD, extensive pairing between a miRNA and a trigger-RNA sequence is thought to drive a conformational change in the argonaute/miRNA/trigger-RNA complex, allowing ZSWIM8 to promote miRNA and argonaute degradation^25,26,36^. This, in turn, is predicted to result in upregulated translation of other miRNA target mRNAs^27^. Comparison of EBAX-1 and ZSWIM8 Alphafold structures^37^, using only high-confidence predicted regions (predicted local distance difference test, pLDDT > 70; for EBAX-1: 993/1717 residues and for ZSWIM8: 927/1837 residues), reveals markedly overlapping 3D alignment (root mean square deviation = 1.112 Å over 399 pruned Cα pairs) (**Figure 5b**), suggesting functional conservation. Indeed, previous studies demonstrated that, like ZSWIM8, EBAX-1 can regulate levels of certain miRNAs^38,39^. To test whether EBAX-1 regulates TDMD to control linker cell death, we first sought to determine if ZSWIM8 can functionally substitute for EBAX-1 in this process. We found that a *mig-24p::Zswim8* cDNA transgene partially, but significantly, restores linker cell death to *ebax-1(tm2321)* mutants (**Figure 5c**), suggesting that EBAX-1 and ZSWIM8 likely carry out similar biochemical functions.

**Figure 5.**
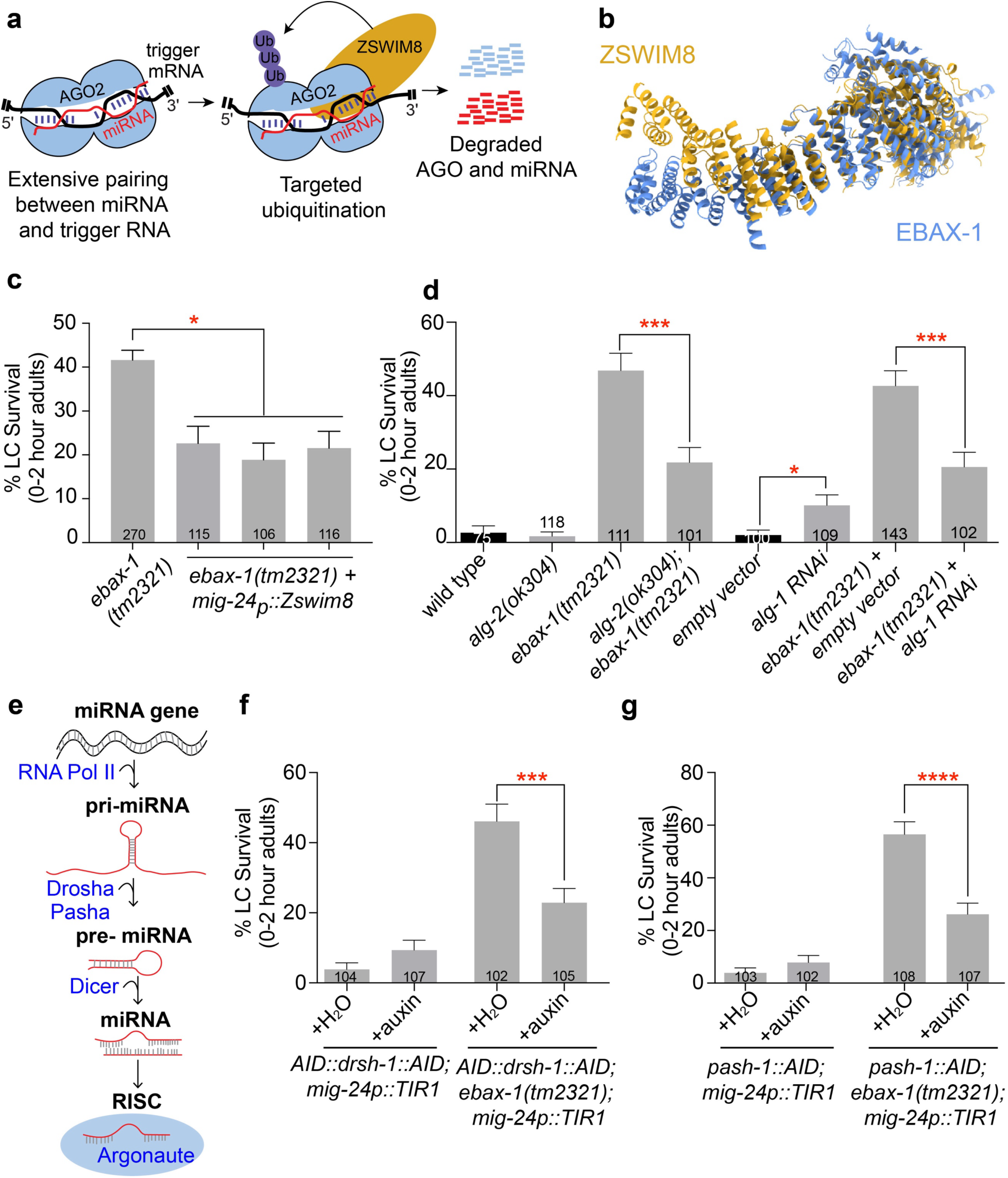
EBAX-1 promotes LCD through its conserved function in target directed miRNA degradation. **(a)** Schematic of target directed miRNA degradation. Adapted from Shi et al., 2020^26^. **(b)** Superposition of *C. elegans* EBAX-1 (blue) and human ZSWIM8 (gold) AlphaFold models. Low-confidence regions (pLDDT < 70) are hidden. **(c)** Linker cell survival in *ebax-1(tm2321)* and *ebax-1(tm2321)* mutant animals expressing human *Zswim8* cDNA in the linker cell (*mig24p::Zswim8*). Three independent lines were tested. From left to right: (*) *p* = 0.0017, 0.0002, and 0.0143; Fisher’s exact test. **(d)** Linker cell survival in indicated genotypes. From left to right: (***) *p* = 0.0002, (*) *p* = 0.02, (***) *p* = 0.0003; Fisher’s exact test. **(e)** miRNA biogenesis pathway. Adapted from Dexheimer et al., 2020^42^. **(f)** Linker cell survival in wild type and *ebax-1(tm2321)* mutant animals that contain auxin-inducible degrons at the *drsh-1* locus, *AID::drsh-1::AID*, and express linker-cell specific TIR1, *mig-24p::TIR1*. Animals were raised on auxin, to deplete DRSH-1 from the linker cell, and on water, as a control. (***) *p* = 0.0005, Fisher’s exact test. **(g)** As in **(f)** but for *pash-1*. (****) *p* < 0.0001, Fisher’s exact test. Bar graph data are plotted as mean ± standard error of the proportion. Number of animals scored listed inside box.

During TDMD, only argonaute proteins associated with miRNAs bound to specific trigger-RNAs are degraded by ZSWIM8 (**Figure 5a**)^27^. Consistent with this, we do not observe gross changes in the levels of ALG-1 or ALG-2, key miRNA-associated *C. elegans* argonaute proteins^40^, during linker cell death in wild-type or in *ebax-1(tm2321)* mutant animals (**Supplementary Figures 4 and 5**). Nonetheless, we found significant suppression of aberrant linker cell survival in *alg-2(ok304); ebax-1(tm2321)* double mutants compared to *ebax-1(tm2321)* single mutants alone (**Figure 5d**). Reducing *alg-1* function using RNAi has a similar effect (**Figure 5D**). Together, these findings are consistent with the notion that TDMD mediated by EBAX-1 promotes linker cell death. Combining *agl-2(ok304)* and *alg-1(RNAi)* does not further suppress linker cell survival in *ebax-1* mutants (**Supplementary Fig. 6**), suggesting that ALG-1 and ALG-2 associate with at least some of the same miRNAs during linker cell death.

Drosha/DRSH-1 and Pasha/PASH-1 are enzymes required for cleavage of primary miRNA transcripts into precursor miRNAs that are subsequently processed by Dicer into mature miRNAs^41^ (**Figure 5e**). To more directly test the role of miRNAs in linker cell death, we crossed published AID-tagged alleles of *drsh-1* and *pash-1*^42^ into wild type or *ebax-1* mutant strains carrying a *mig-24p::TIR1::mRuby* transgene expressing linker-cell specific TIR1. While depletion of DRSH-1 or PASH-1 from wild-type linker cells by exposure to auxin does not affect linker cell death, removal of either protein in *ebax-1* mutants restores linker cell death (**Figure 5f,g**).

Our findings are, therefore, consistent with the notion that EBAX-1 functions in TDMD during linker cell death to target ALG-1 and ALG-2 miRNA complexes bound to specific trigger RNAs for degradation.

### EBAX-1 works through *mir-35* family miRNAs to promote linker cell death

We previously described a role for *let-7* miRNA, acting as part of a developmental timing pathway, in linker cell death activation^14^. Furthermore, ALG-1 has been reported to function with *let-7* miRNA in other contexts^40,43–45^. Supporting a role for ALG-1/*let-7* miRNA interactions during linker cell death, *alg-1(RNAi)* causes a mild block in linker cell death (**Figure 5d**). Nonetheless, *let-7* is unlikely to be an EBAX-1 TDMD target during linker cell death, as *alg-1(RNAi)* suppresses ectopic linker cell survival in *ebax-1(tm2321)* mutants (**Figure 5d**) and loss of *let-7* blocks linker cell death instead of promoting it^14^. We conclude that although *alg-1* normally promotes linker cell death through its interaction with *let-*7, it also likely functions with a different miRNA to block linker cell demise.

What might this miRNA be? The *mir-35* family of miRNAs is a known TDMD target of EBAX-1 in early larval development^38,39^. This miRNA family comprises eight closely related miRNAs (*mir-35-42*) that share a conserved seed sequence, which mediates post-transcriptional repression of target mRNAs (5’CACCGGG3’; **Figure 6a**)^46^. To assess whether the *mir-35* miRNA family regulates linker cell death, we scored linker cell survival in animals homozygous for the *nDf50* single genomic deletion or *nDf50 nDf49* double deletions, which remove *mir-35-41* and *mir-35-42*, respectively. While either deletion alone does not perturb linker cell death, each deletion significantly suppresses inappropriate linker cell survival in *ebax-1* mutants (**Figure 6b**). Thus, *mir-35* family miRNAs act downstream of EBAX-1 and are likely EBAX-1 targets during linker cell death. Furthermore, over-expression of *mir-35-41* genomic sequences in wild-type linker cells causes significant inappropriate linker cell survival (**Figure 6c**), suggesting that *mir-35* family miRNAs block linker cell death. Together our results indicate that EBAX-1-dependent degradation of *mir-35* family miRNAs promotes linker cell demise.

**Figure 6.**
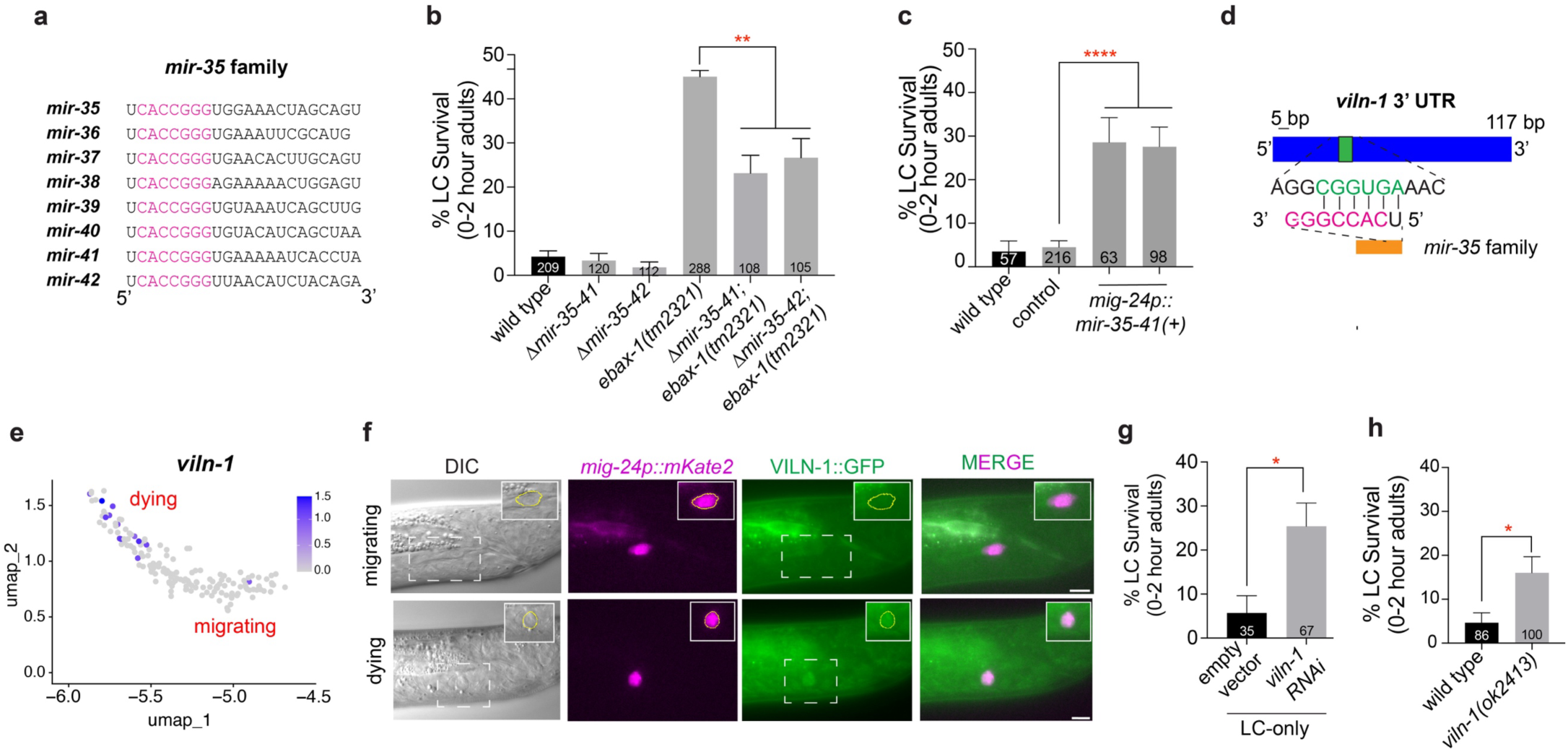
EBAX-1 inhibits *mir-35* family miRNAs allowing for target gene upregulation. **(a)** The *mir-35* miRNA family consists of 8 miRNAs, *mir-35*, *36, 37, 38, 39, 40, 41* and *42*, that share a common seed sequence that recognizes target mRNAs, CACCGGG (pink). **(b)** Linker cell survival in indicated genotypes. Δ*mir-35-41*, *nDf50*. Δ*mir-35-42*, *nDf50 nDf49*. From left to right: (**) *p* = 0.0002, 0.0064; Fisher’s exact test. **(c)** Linker cell survival in wild type animals expressing *mig-24p::mir-35-41* transgene. Wild type non-transgenic siblings were used as the control. Wild type, parental wild type strain used to generate transgenic *mig-24p::mir-35-41* animals. Two independent lines were tested. (****) *p* < 0.0001, Fisher’s exact test. **(d)** Schematic of the 3’ untranslated region (UTR) of *viln-1* showing the predicted seed sequence (green) that is recognized by the *mir-35* miRNA family seed sequence (pink). **(e)** *viln-1* expression in migrating and dying linker cells transcriptionally profiled using single-cell RNA sequencing. **(f)** Representative DIC and fluorescent micrographs of VILN-1::GFP expression in a linker cell finishing migration (top) and a dying linker cell (bottom). *mig-24p::mKate2* was used as a linker cell marker. Inset, linker cell. Scale bar, 5 μm. **(g)** Linker cell survival in *him-8(e1489); rde-1(ne219) qIs56* mutants expressing *mig-24p::rde-1* cDNA treated with empty vector or *viln-1* RNAi. (*) *p* = 0.0163, Fisher’s exact test. **(h)** Linker cell survival in indicated genotypes. (*) *p* = 0.0164, Fisher’s exact test. Bar graph data are plotted as mean ± standard error of the proportion. Number of animals scored listed inside box.

### *viln-1* and *lin-29* predicted *mir-35* family miRNA targets are required for linker cell death

Our findings suggest that pro-death mRNAs, whose translation is blocked by *mir-35* family miRNAs, become activated through TDMD to promote linker cell death. To test this idea, we sought to identify such *mir-35* family target mRNAs. We first examined the 3’ untranslated regions (UTR) of known linker cell death genes^18^ for sites complementary to the *mir-35* family seed sequence using TargetScanWorm^47,48^ (**Figure 6a**). We noted that the 3’ UTR positions 327-333 of the *lin-29* mRNA, encoding a Zn-finger transcription factor required for linker cell death^14^, are complementary to positions 1-7 of *mir-35* family microRNAs (**Supplementary Fig. 7a**). Furthermore, the site is conserved in the related nematodes *C. remanei, C. briggsae*, and *C. brenneri* (**Supplementary Fig. 7b**). However, previous studies from our group are consistent with LIN-29, a component of the developmental timing pathway, acting upstream of HSF-1^19^ (**Figure 4d**). Furthermore, *lin-29* mRNA accumulation is not induced at the onset of linker cell death and is expressed in both migrating and dying linker cells^14^ (**Supplementary Fig. 7d**). Thus, while *lin-29* mRNA is a likely target of *mir-35* family inhibition, which may be relieved by EBAX-1, this activity does not account for our finding that EBAX-1 lies genetically downstream of HSF-1 (**Figure 4**).

We recently generated a developmental transcriptome tracking the linker cell as it transitions from migration to death, identifying 297 transcripts upregulated in dying cells^49^. We also searched the predicted 3’ UTRs of these transcripts for complementarity to the *mir-35* family seed sequence and identified seven genes whose mRNAs are upregulated in the dying linker cell and are potential *mir-35* family targets. Among these is *viln-1* mRNA, encoding the actin-binding cytoskeletal protein villin. Positions 34-39 of the *viln-1* 3’ UTR are complementary to residues 1-6 of all *mir-35* family miRNAs (**Figure 6d**) and position 31 of the 3’ UTR is complementary to position 9 of all, but one, *mir-35* family members. Unlike *lin-29*, *viln-1* mRNA expression is upregulated in dying linker cells according to our RNAseq data (**Figure 6e**). Furthermore, expression of an endogenous *viln-1::gfp* gene fusion we generated by CRISPR/Cas9 is barely detected in the migrating linker cell but easily observed in the dying linker cell (**Figure 6f, Supplementary Fig. 8** n=27 animals). Importantly, *viln-1(ok413)* mutants, as well as animals exposed to linker cell specific *viln-1* RNAi, exhibit significant linker-cell-survival (**Figure 6g,h**). These observations are consistent with the notion that *viln-1* mRNA may accumulate, at least in part, through EBAX-1-mediated degradation of *mir-35* family miRNAs to promote linker cell death.

## DISCUSSION

The data presented here uncover a novel role for TDMD in promoting programmed cell death during animal development. Our findings suggest a model for the control of linker cell death in which EBAX-1, acting as a Cullin-2-based E3 ubiquitin ligase component, degrades the argonautes ALG-1 and ALG-2 on specific *mir-35* miRNA-mRNA complexes. This leads to the elimination of *mir-35* family miRNAs by target-directed miRNA degradation, which, in turn, allows accumulation of *viln-1* villin, required for LCD (**Figure 7**). Missing from this model is the identity of the trigger-RNA responsible for *mir-35* family miRNA degradation. One intriguing possibility is that *lin-29* mRNA is this molecule. Indeed, alignment of *lin-29* 3’ UTR and *mir-35* miRNA sequences reveals additional binding potential beyond the seed sequence in a pattern reminiscent of previously validated trigger-RNAs^36^ (**Supplementary Fig. 7c**). We identified two potential binding configurations (**Supplementary Fig. 7c**) that could increase the miRNA-mRNA binding constant, increasing the likelihood of triggering TDMD.

**Figure 7.**
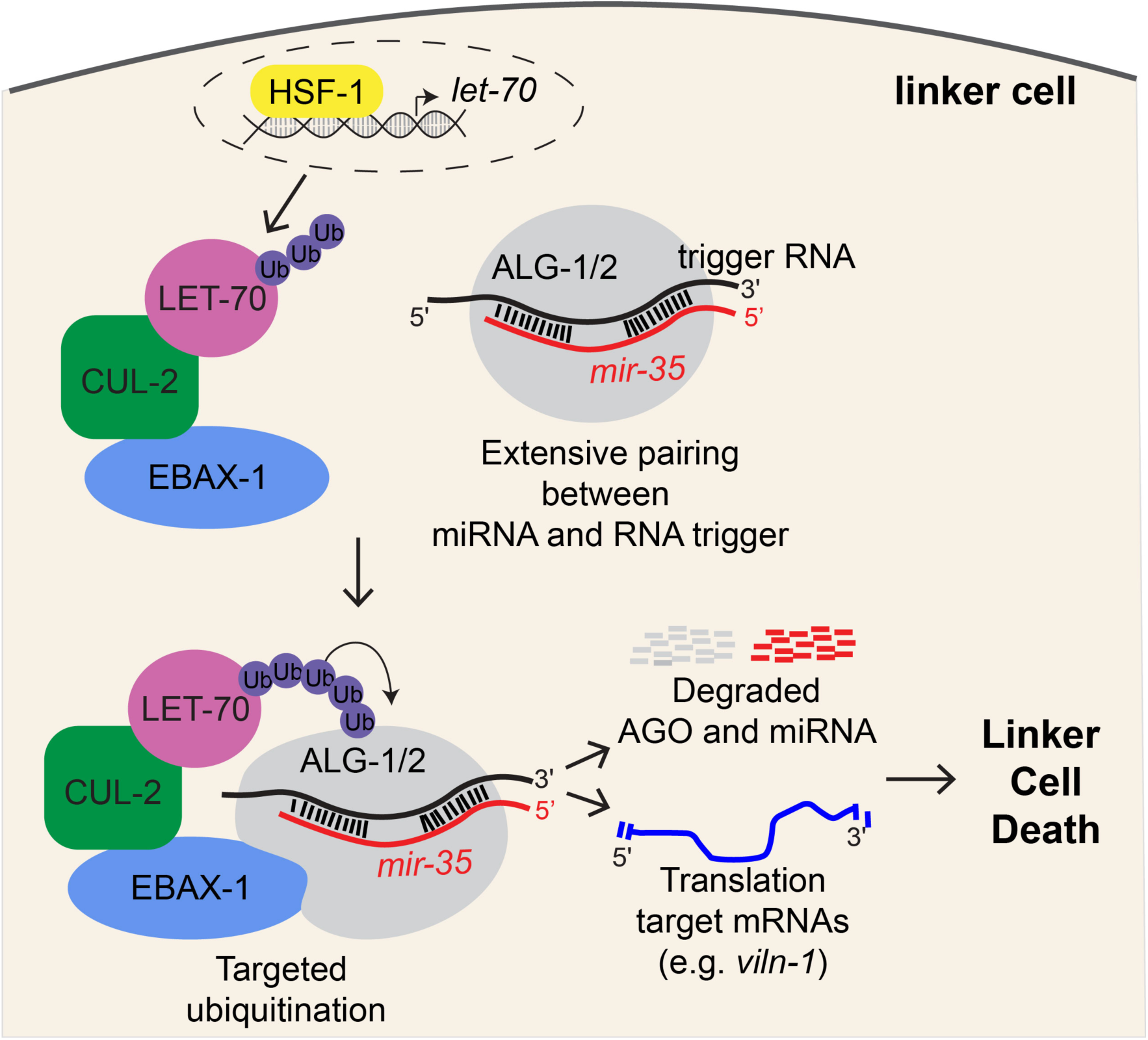
A model for the role of EBAX-1 and TDMD in linker cell death. HSF-1 activates transcription of the E2 ubiquitin conjugating enzyme LET-70. LET-70 interacts with the Cul2-based E3 ligase complex containing EBAX-1 as the substrate recognition domain. Extensive pairing between a *mir-35* family miRNA and a yet unknown trigger RNA sequence causes a conformational change in the associated argonautes, ALG-1/2, which is then recognized by EBAX-1. ALG-1/2 are then slated for degradation by the proteasome along with the associated miRNA. Upon miRNA degradation, target mRNAs that are normally repressed by the *mir-35* family, such as *viln-1*, have increased translation and promote cell death.

### Functional relevance of TDMD to animal development

The biochemical mechanisms mediating TDMD have been intensely investigated^27,50^. Extensive complementarity between a miRNA and its trigger RNA is thought to alter the conformation of argonaute proteins bound to this miRNA-RNA complex, allowing recognition by the ZSWIM8/EBAX-1-CUL-2 ubiquitin ligase complex, which mediates subsequent degradation of the argonaute protein and its bound miRNA^25–27^. The physiological contexts in which TDMD functions, however, are only beginning to be defined. In *Drosophila*, the ZSWIM8 homolog *Dora* mediates degradation of multiple miRNAs required for embryonic viability, including *miR-310*, whose loss promotes cuticle development^51^. In mice, *ZSWIM8* knockout increases expression of many miRNAs and animals display a range of developmental abnormalities, including heart and lung defects, a small body size, and perinatal lethality^52,53^. Loss of two miRNAs, *miR-322* and *miR-503*, suppresses the body size defects of *ZSWIM8* knockout mice^52^, implicating TDMD in body size control. In *C. elegans*, a long non-coding RNA, *tts-*2, was recently shown to trigger EBAX-1-dependent decay of all *mir-35* family members at the embryo-to-larval transition^38,54^; and in the nematode *Pristionchus pacificus*, EBAX-/ZSWIM8 destabilizes *mir-2235a/mir-35* family miRNAs to mediate the transgenerational inheritance of a predatory trait^55^. Our findings extend these studies, demonstrating that TDMD can act not only as a homeostatic mechanism for miRNA turnover during embryonic development, but as an active molecular switch that drives an irreversible developmental event – cell death.

### Is TDMD a bridge between apoptosis and LCD?

Our findings raise the possibility that selective miRNA degradation may be a general strategy for committing cells to death. Supporting this idea, *mir-35* family miRNAs were previously shown to control apoptosis in *C. elegans* by repressing *egl-*1/BH3-only mRNA, encoding an activator of the core apoptotic pathway^56^. Furthermore, in germ cells that undergo DNA-damage induced apoptosis, *mir-35* family miRNAs inhibit cell death by repressing expression of the MAPK signaling regulator NDK-1^57^. More broadly, a recent report suggests that although caspases are not required for *C. elegans* linker cell death, they nonetheless play an important role in assuring the fidelity of its execution^58^. Together, these various observations raise the possibility that although the details may differ, underlying connections between developmental cell death programs exist, which may reflect a common evolutionary origin.

### Additional miRNAs and EBAX-1 Substrates involved in LCD

Our data suggest that the linker cell death TDMD pathway we unveiled is likely to be more extensive. Our finding that a putative *ebax-1* null mutant does not fully eliminate linker cell death (**Figure 1D, E**), for example, suggests that other E3 complexes may drive LCD; a result supported by our finding that loss of *btbd-*2, another E3 component, enhances linker cell survival in *ebax-1* mutants^19^ (**Figure 1F)**. Likewise, our data suggest that *mir-35* family miRNAs may not be the only targets of EBAX-1, as loss of this miRNA family does not fully restore LCD to *ebax-1* mutants (**Figure 6B**). Indeed, previous work showed that EBAX-1 controls the stability of numerous miRNAs throughout *C. elegans* development and in adulthood^26,39^. The partial defects observed in *viln-1* mutants (**Figure 6F, G)**, support the idea that additional *mir-35* family mRNA targets may also exist. Degradation of multiple miRNA families and activation of their associated target mRNAs may act in concert to enable a broad transcriptional reset that drives linker cell death. Thus, TDMD could serve as a combinatorial mechanism that synchronously releases multiple mRNA programs necessary for execution of LCD. Defining these additional substrates will be key to understanding how the ubiquitin proteasome system integrates post-transcriptional and proteolytic control to orchestrate non-apoptotic cell death.

## METHODS

### Strains

All *C. elegans* strains used in this study were cultured using standard methods^59^ and grown at 20°C, unless indicated otherwise. Wild-type animals were of the Bristol N2 strain.

Most strains harbor the *him-5(e1490)* mutation that generates a high incidence of male progeny, as well as one of two integrated linker cell markers, *qIs56[lag-2p::GFP]* or *nsIs650[mig-24p::mKate2]*. A complete list of strains used and generated in this study is found in **Table S1**.

### CRISPR/Cas9 Genome editing

CRISPR/Cas9 genome editing was performed by injection of Cas9, CRISPR RNAs (crRNAs), and trans activating crRNAs (tracrRNAs) from Integrated DNA Technologies (IDT) as described in Dokshin et al., 2018^30^ and Eroglu et al., 2023^60^. Generation of the *ebax-1(ns1019)* deletion allele was performed by use of two crRNAs and a single-stranded oligodeoxynucleotide (ssODN) that was purchased as an Ultramer DNA Oligo from IDT. The *ebax-1(ns1019)* deletion was originally made in the *let-70(ns770)* mutant strain because *let-70* and *ebax-1* are tightly linked on Chr IV. *ebax-1(ns1019)* was also separately generated precisely in a wild-type background.

Generation of the *viln-1* knock in reporter was performed by a single crRNA and an ssODN repair donor template. The ssODN, including the desired insertion sequence flanked by homology arms, was prepared by enzymatic digestion of a PCR product into single stranded DNA as described in Eroglu et al. 2023^60^. Sequences of the oligonucleotides used in this study are listed in **Table S2**. The following alleles were generated using CRISPR/Cas9 by SUNY Biotech:

1. *ebax-1(syb2784[ebax-1::GFP]),* C terminal tagging of EBAX-1 with GFP.
2. *ebax-1(syb7218[ebax-1::3xGAS::mNeonGreen::AID*], C terminal tagging of EBAX-1 with linker::mNeonGreen::AID.
3. *tag-30(syb7669),* precise 901 bp deletion within the *tag-30* coding sequence that removes exons 2 through 5. TAG-30 is the *C. elegans* homolog of human *BTBD-2*; throughout this manuscript we refer to *tag-30* as *btbd-2. tag-30(syb7669)* deletion was made separately in both the wild-type and in *ebax-1(tm2321)* mutants because *ebax-1* and *tag-30* are tightly linked on Chr IV.
4. *alg-2(syb8021[alg-2::GFP-PEST]),* C terminal tagging of ALG-2 with GFP-PEST.

### Plasmid Construction

Most plasmids used in this study were made using Gibson assembly^61^. The *mig-24p::ebax-1* Δ*BC-box*, *mig-24p::ebax-1* Δ*SWIM*, and *mig-24p::ebax-1* Δ*A* plasmids were made using restriction free cloning^62^. The *mig-24p::ebax-1* Δ*Cul2-box* was made using the Q5 Site-Directed Mutagenesis Kit (New England Biolabs). A complete list of all plasmids used in this study along with details about their construction are found in in **Table S3.**

### Germline transformation and integration

Transgenic lines were generated by injection of plasmid DNA mixes into the hermaphrodite gonad as previously described^63^. The *ebax-1* rescuing fosmid was injected as a simple extrachromosomal array at 5 ng/μL with *odr-1p::RFP* (20 ng/μL) and *lag-2p::mCherry* (20 ng/μL) as co-injection markers, and pBlueScript (55 ng/μL) as filler DNA to reach a minimum DNA injection concentration of 100 ng/μL. The integrated array, *nsIs1045[mig-24p::TIR1::mRuby + unc-122p::mCherry]*, was generated by exposing animals to 33.4 μg/mL trioxsalen (Sigma T6137) and UV irradiation using a Stratagene Stratalinker UV 2400 Crosslinker (360 μJ/cm^2)^ as previously described^64^. *nsIs1045* animals were outcrossed at least 4 times before being used in experiments.

### Linker Cell Survival Assays

Linker cell death was scored as previously described^20^. Briefly, gravid hermaphrodites were treated with alkaline bleach to isolate eggs, which were allowed to hatch overnight in M9 buffer. Synchronized L1 larvae were plated onto NGM plates seeded with OP50 and maintained at 20°C. Approximately two days after plating, when most animals reached the L4 stage, males exhibiting a fully retracted tail tip with rays visible beneath the unshed L4 cuticle (L4-molt stage) were transferred to fresh plates. Two hours later, newly molted adults were mounted on 2% agarose-M9 pads, anesthetized with 25 mM sodium azide and examined using Nomarski optics and widefield fluorescence on a Zeiss AxioScope A1 microscope (63x oil objective). The linker cell was identified by its location and morphology, as well as by green fluorescence from reporter transgenes. A linker cell was scored as surviving if the nucleus appeared circular with an intact nucleolus, the cell shape was not rounded and refractile, and if the cell had not shed any large blebs. All other cells were classified as dead or dying.

To quantify linker cell death in 1-day old adults, synchronized L4 molt males were transferred to new plates and scored 24 hours later using the same criteria described above.

For the rescue experiments, extra-chromosomal arrays carrying the rescuing construct included a bright *unc-122p::mCherry* co-injection marker. Males expressing *mCherry* fluorescence were pre-selected under a fluorescence dissection scope prior to scoring at the L4 molt stage. To assess rescue, *mCherry*-positive (transgenic) and *mCherry*-negative (non-transgenic) siblings from the same strain were scored and compared in parallel.

For scoring *ebax-1(ns1019) let-70(ns770)* mutants, as well as animals harboring the *nDf50* and *nDf49 nDf50* alleles, animals were maintained as heterozygotes using balancers for Chromosomes IV (*nT1*) and II (*mIn1*), respectively, and balancer negative homozygous animals were isolated before scoring.

### RNAi experiments

RNA interference (RNAi) experiments were performed by feeding, as previously described^65,66^. Briefly, bleached embryos from gravid hermaphrodites were synchronized at the L1 stage by leaving them overnight in M9. Synchronized L1 larvae were placed on NGM plates with 1 mM IPTG and 25 μg/mL carbenicillin and coated with bacteria carrying the desired RNAi clone or empty vector RNAi control plasmids. Worms were grown for approximately 48 hours at 20°C and scored for linker cell death, as described above. RNAi clones were isolated from the Ahringer feeding library^66^ and each clone’s sequence was confirmed before use with Sanger sequencing. RNAi targeting GFP served as positive control, and animals were scored for efficient GFP knockdown (≥60%) before experimental scoring. For linker cell specific RNAi, we used a strain deficient for RNAi (*rde-1(ne219)*) but also expressing linker cell-specific *rde-1* cDNA, rendering the RNAi machinery functional specifically in linker cells, as previously described^19^.

### Temporally controlled linker cell-specific protein degradation

To degrade EBAX-1, DRSH-1 and PASH-1 in the linker cell, we used the auxin-inducible degradation system^33^, in which animal exposure to a synthetic auxin analog, K-NAA, results in the ubiquitination and subsequent degradation of auxin-inducible degron (AID)-tagged proteins in cells that also express TIR1, the substrate recognition component of the E3 ubiquitin ligase complex^32,33^. Strains carrying *ebax-1::mNeonGreen::AID* [*ebax-1(syb7218)*], *AID::drsh-1::AID* [*drsh-1(luc82)*], and *pash-1::AID* [*pash-1(luc71)*] were crossed into a strain expressing TIR1 ubiquitously (*eft-3p::TIR1::mRuby*) and/or a strain expressing TIR1 specifically in the linker cell (*nsIs1045[mig-24p::TIR1]*). K-NAA, 1-napthaleneacetic Acid Potassium Salt (PhytoTech Labs #N610), was dissolved in sterile H2O to prepare a 200 mM stock solution. OP50-seeded NGM plates were coated with K-NAA for a final concentration of 4 mM. Synchronized populations of L1 animals were transferred onto K-NAA plates and grown at 20°C until animals reached the late L4 larval stage and could be scored for LCD. Age-matched animals placed on OP50-seeded NGM plates coated with sterile H2O were used as controls.

For the EBAX-1 time of action experiment (**Figure 2f,g, Supplementary Fig. 2**), synchronized populations of L1 animals were transferred onto K-NAA plates or H2O plates and grown at 20°C. Twenty four later, animals were thoroughly washed in M9 and transferred onto H2O plates or K-NAA plates, respectively, and raised until animals were ready to be scored for LCD.

### Electron microscopy

Young adult (0-2 hours post L4 molt) *him-5(e1490) qIs56[lag-2p::GFP]* males with dying linker cells, and young adult and adult (24 hours post L4 molt) *ebax-1(tm2321); him-5(e1490) qIs56[lag-2p::GFP]* males with surviving linker cells were imaged using a Zeiss compound microscope (Axio Imager M2, 63x objective) to measure the relative location of the linker cell within the worm using ImageJ software^67^. Animals were then fixed, stained, embedded in resin, and sectioned using standard methods^68^. Images were acquired using a Titan Themis 200 kV Transmission electron microscope with Cs Image Corrector. Image processing and analysis were performed using ImageJ and IMOD software.

### Microscopy

Animals were mounted on 2% agarose pads made in M9 medium and immobilized with 25 mM or 250 mM sodium azide for widefield or confocal microscopy, respectively.

Differential Interference Contrast (DIC) and widefield fluorescence micrographs (images in Figures 1B, 1C, 6E, S8) and widefield fluorescence z-stacks (each ∼1-3 μm thick, images in Figure 2B), were acquired using a Zeiss compound microscope (Axio Imager M2) using a 63x/1.4 NA oil objective controlled by MicroManager software (v1.4.22)^69^.

Confocal z-stack images (each ∼0.1 - 0.35 μm thick) were acquired using a Zeiss LSM900 inverted laser scanning confocal microscope controlled by ZenBlue software with a 63x/1.4 NA oil objective under standard confocal mode (images in **Figure 2e**, **Supplementary Fig. 4**) or Airyscan super resolution mode (images in **Supplementary Figures 2 and 5**). Images acquired in the Airyscan super resolution mode were deconvoluted using the Airyscan Processing algorithm on the ZenBlue software before being further analyzed in Fiji^67^. Maximum and sum-projections of all z-stack images presented were prepared using Fiji^67^, and figures were prepared using Adobe Illustrator.

Z-stack images of EBAX-1::GFP localization in the linker cell (images in **Figure 4e**) were acquired using a Leica DMi8 inverted microscope with InstantSIM (iSIM) real-time super resolution with the 63x/1.3 NA glycerol objective controlled by VisiView acquisition software.

### Quantification of EBAX-1::GFP Fluorescence

Fluorescence intensity measurements were performed on z-stack images acquired with the iSIM microscope using identical laser and acquisition settings for all samples. To correct for day-to-day variation in illumination and detector sensitivity, a blank calibration image was taken each imaging session using the same region of the field of view, laser power, and exposure time as the experimental images. All image analysis were performed in Fiji^67^. Calibration images were processed by subtracting the minimum pixel value, converting the image to 32 bit, and then dividing by the new maximum pixel value to generate a normalized calibration image. For each sample z-stack, a sum-intensity projection of three GFP z-slices (∼1.0 μm interval) encompassing the linker cell was generated. To correct for uneven illumination across the field of view, each sum-projection image was divided by the corresponding normalized calibration image using Fiji’s

*Image Calculator* function. Pixel intensity measurements were then performed on the resulting normalized sum-projection image. Regions of interest (ROI) were manually drawn around the entire cell and the nucleus, and the mean fluorescence intensities were measured (𝐼_𝑐𝑒𝑙𝑙_ and 𝐼_𝑛𝑢𝑐𝑙𝑒𝑢𝑠_, respectively). An ROI was also drawn around a nearby background region to obtain the background intensity (𝐼_𝑏(_). Background-subtracted intensities were defined as:

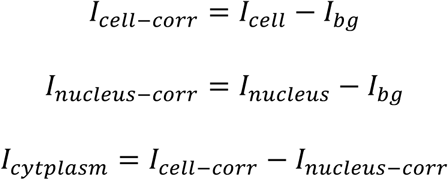

The nuclear to cytoplasmic fluorescence ratio was then calculated as:

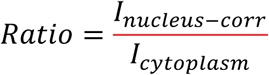

All quantifications were performed on images acquired from wild type (empty vector) and *let-70 RNAi* treated animals imaged on the same day to minimize variability due to laser fluctuations.

### Protein structure modelling and superposition

AlphaFold2 models for human ZSWIM8 (UniProt A7E2V4) and *C. elegans* EBAX-1 (UniProt Q21875) were downloaded from the AlphaFold Protein Structure Database as PDB files^37^. In these files, per-residue pLDDT (predicted Local Distance Difference Test) confidence values are stored in the B-factor column. Models were opened in UCSF ChimeraX(version 1.10.1)^70^. To focus on well-predicted regions of the proteins, residues with a pLDDT < 70 were excluded from visualization (selected by B-factor and hidden), so only high-confidence residues (pLDDT > 70) are shown. Structures were superimposed using MatchMaker with default settings (Needleman-Wunsch sequence alignment, BLOSUM-62, secondary-structure fraction 0.3; gap-open 18/18/6, gap-extend 1), and Cα Root Mean Square Deviation (RMSD) was computed after automatic pruning of outliers.

### Target ScanWorm Analysis

Predicted *C. elegans* miRNA target sites were identified using TargetScanWorm release 6.2^47,48^ (available at https://www.targetscan.org/worm_52/). The 3’ untranslated regions (3’UTRs) of known linker cell death genes and genes transcriptionally upregulated in dying linker cells^49^ were searched for conserved and non-conserved sites complementary to the *mir-35* family seed sequence (5’CACCGGG-3’). Predicted sites were classified according to canonical TargetScan nomenclature (8-mer-1A, 7-mer-1A, 6-mer-1A, etc), which describe the degree of base pairing between the miRNA seed and target site. The PCT value (probability of preferential conservation) provided by TargetScan was used as an indicator of evolutionary conservation across *Caenorhabditis* species.

### Quantification and Statistical Analysis

Statistical analysis was performed using Excel and GraphPad Prism software (v. 10). Statistical parameters including mean ± standard error of the proportion, mean ± standard error of the mean (SEM) and N are reported in the main text, figures, and figure legends. Data is judged to be statistically significant when *p* < 0.05 by Fisher’s exact test or student’s t-test, where appropriate. Graphs were prepared using GraphPad Prism.

### Genetic Data Analysis

When comparing a transgenic line to a parental strain, a minimum of two independent lines were scored. Approximately 100 animals from each of these lines were examined for linker cell defects and compared to approximately 100 non-transgenic siblings of the same strain using a Fisher’s exact test to determine significance. For ease of presentation, we pooled the lines of non-transgenic siblings from a given experiment and displayed them on the relevant figure using mean ± SEM (N=2 lines).

## DATA AVAILABILITY

All reagents are available from the corresponding author upon request.

## ACKNOWLEDGMENTS

We thank: Katherine McJunkin for useful discussions, *mir-35* family deletion strains and the *mir-35-41* rescue plasmid; Luisa Cochella for useful discussions; Mohammed Alam for generating the single *ebax-1(ju699)* and *ebax-1(tm2321)* mutant strains and the *ebax-1* genomic rescue strain; Lena Kutscher for analysis of the VC40125 million mutation strain; Alison J. North, Christina Pyrgaki, Ved Sharma and Tao Tong from the Rockefeller University’s Bio-Imaging Resource Center, RRID:SCR_017791, for help with iSIM imaging and image analysis; Chris Hammel for the GFP-PEST plasmid; members of the Shaham lab for useful discussions and reagents; the Imaging and the Electron Microscope Imaging Facility of CUNY Advanced Science Research Center for instrument use and technical assistance. iSIM imaging was performed at the Rockefeller University’s Bio-Imaging Resource Center, RRID:SCR_017791. Some strains were provided by the CGC, which is funded by the NIH Office of Research Infrastructure Programs (P40OD010440). This work was supported by the Harvey L. Karp Postdoctoral Fellowship at Rockefeller University to L. B. H., NIH F32GM145036 to L.B.H., and NIH grants R35NS105094 and R01HD103610 to S. S.

## AUTHOR CONTRIBUTIONS

Conceptualization: L.B.H. and S.S.; Methodology: L.B.H .and S.S.; Formal Analysis: L.B.H.; Investigation: L.B.H., O.Y., and Y.L.; Writing – original draft: L.B.H.; Writing – reviewing and editing: L.B.H. and S.S.; Funding acquisition: L.B.H. and S.S.; Resources: L.B.H., O.Y. and S.S.; Visualization: L.B.H., O.Y. and S.S.; Supervision: S.S.

## MATERIALS AND CORRESPONDENCE

Further information and requests for resources and reagents should be directed to the lead contact, Shai Shaham.

## SUPPLEMENTAL INFORMATION

Supplementary Information: Figures S1 – S8, Tables S1, S2, S3.

## REFERENCES

1. Fuchs, Y. & Steller, H. Live to die another way: modes of programmed cell death and the signals emanating from dying cells. Nat Rev Mol Cell Biol 16, 329–44 (2015).

2. Ghose, P. & Shaham, S. Cell death in animal development. Development 147, dev191882 (2020).

3. Fuchs, Y. & Steller, H. Programmed cell death in animal development and disease. Cell 147, 742–58 (2011).

4. Horvitz, H. R. NOBEL LECTURE: Worms, Life and Death. Biosci. Rep. 23, 239–303 (2003).

5. Thornberry, N. A. & Lazebnik, Y. Caspases: Enemies Within. Science 281, 1312–1316 (1998).

6. Kerr, J. F. R., Wyllie, A. H. & Currie, A. R. Apoptosis: A Basic Biological Phenomenon with Wideranging Implications in Tissue Kinetics. Br. J. Cancer 26, 239–257 (1972).

7. Xu, D. et al. Genetic control of programmed cell death (apoptosis) in Drosophila. Fly 3, 78–90 (2009).

8. Honarpour, N. et al. Adult Apaf-1-deficient mice exhibit male infertility. Dev Biol 218, 248– 58 (2000).

9. Kuida, K. et al. Reduced apoptosis and cytochrome c-mediated caspase activation in mice lacking caspase 9. Cell 94, 325–37 (1998).

10. Ke, F. F. S. et al. Embryogenesis and Adult Life in the Absence of Intrinsic Apoptosis Effectors BAX, BAK, and BOK. Cell 173, 1217–1230 e17 (2018).

11. Hanahan, D. & Weinberg, R. A. Hallmarks of Cancer: The Next Generation. Cell 144, 646– 674 (2011).

12. Dhani, S., Zhao, Y. & Zhivotovsky, B. A long way to go: caspase inhibitors in clinical use. Cell Death Dis. 12, 949 (2021).

13. Li, K., Delft, M. F. van & Dewson, G. Too much death can kill you: inhibiting intrinsic apoptosis to treat disease. EMBO J. 40, e107341 (2021).

14. Abraham, M. C., Lu, Y. & Shaham, S. A morphologically conserved nonapoptotic program promotes linker cell death in Caenorhabditis elegans. Dev Cell 12, 73–86 (2007).

15. Kimble, J. & Hirsh, D. The postembryonic cell lineages of the hermaphrodite and male gonads in Caenorhabditis elegans. Dev Biol 70, 396–417 (1979).

16. Kutscher, L. M., Keil, W. & Shaham, S. RAB-35 and ARF-6 GTPases Mediate Engulfment and Clearance Following Linker Cell-Type Death. Dev Cell 47, 222–238 e6 (2018).

17. Denning, D. P., Hatch, V. & Horvitz, H. R. Both the caspase CSP-1 and a caspase-independent pathway promote programmed cell death in parallel to the canonical pathway for apoptosis in Caenorhabditis elegans. PLoS Genet 9, e1003341 (2013).

18. Horowitz, L. B. & Shaham, S. Apoptotic and Nonapoptotic Cell Death in Caenorhabditis elegans Development. Annu. Rev. Genet. 58, 113–134 (2024).

19. Kinet, M. J. et al. HSF-1 activates the ubiquitin proteasome system to promote non-apoptotic developmental cell death in C. elegans. Elife 5, (2016).

20. Blum, E. S., Abraham, M. C., Yoshimura, S., Lu, Y. & Shaham, S. Control of nonapoptotic developmental cell death in Caenorhabditis elegans by a polyglutamine-repeat protein. Science 335, 970–3 (2012).

21. Kutscher, L. M. & Shaham, S. Non-apoptotic cell death in animal development. Cell Death Differ 24, 1326–1336 (2017).

22. Davies, S. W. et al. Formation of neuronal intranuclear inclusions underlies the neurological dysfunction in mice transgenic for the HD mutation. Cell 90, 537–48 (1997).

23. Turmaine, M. et al. Nonapoptotic neurodegeneration in a transgenic mouse model of Huntington’s disease. Proc Natl Acad Sci U S A 97, 8093–7 (2000).

24. Osterloh, J. M. et al. dSarm/Sarm1 is required for activation of an injury-induced axon death pathway. Science 337, 481–484 (2012).

25. Han, J. et al. A ubiquitin ligase mediates target-directed microRNA decay independently of tailing and trimming. Science 370, (2020).

26. Shi, C. Y. et al. The ZSWIM8 ubiquitin ligase mediates target-directed microRNA degradation. Science 370, (2020).

27. McJunkin, K. & Gottesman, S. What goes up must come down: off switches for regulatory RNAs. Genes Dev. 38, 597–613 (2024).

28. Thompson, O. et al. The million mutation project: A new approach to genetics in Caenorhabditis elegans. Genome Res. 23, 1749–1762 (2013).

29. Wang, Z. et al. The EBAX-type Cullin-RING E3 Ligase and Hsp90 Guard the Protein Quality of the SAX-3/Robo Receptor in Developing Neurons. Neuron 79, 903–916 (2013).

30. Dokshin, G. A., Ghanta, K. S., Piscopo, K. M. & Mello, C. C. Robust Genome Editing with Short Single-Stranded and Long, Partially Single-Stranded DNA Donors in Caenorhabditis elegans. Genetics 210, 781–787 (2018).

31. Ashley, G. E. et al. An expanded auxin-inducible degron toolkit for Caenorhabditis elegans. Genetics 217, iyab006 (2021).

32. Zhang, L., Ward, J. D., Cheng, Z. & Dernburg, A. F. The auxin-inducible degradation (AID) system enables versatile conditional protein depletion in C. elegans. Development 142, 4374– 4384 (2015).

33. Martinez, M. A. Q. et al. Rapid Degradation of Caenorhabditis elegans Proteins at Single-Cell Resolution with a Synthetic Auxin. *G*3: Genes, Genomes, Genet. 10, 267–280 (2020).

34. Mahrour, N. et al. Characterization of Cullin-box Sequences That Direct Recruitment of Cul2-Rbx1 and Cul5-Rbx2 Modules to Elongin BC-based Ubiquitin Ligases*. J. Biol. Chem. 283, 8005–8013 (2008).

35. Makarova, K. S., Aravind, L. & Koonin, E. V. SWIM, a novel Zn-chelating domain present in bacteria, archaea and eukaryotes. Trends Biochem. Sci. 27, 384–386 (2002).

36. Buhagiar, A. F. & Kleaveland, B. To kill a microRNA: emerging concepts in target-directed microRNA degradation. Nucleic Acids Res. 52, 1558–1574 (2024).

37. Jumper, J. et al. Highly accurate protein structure prediction with AlphaFold. Nature 596, 583–589 (2021).

38. Donnelly, B. F. et al. The developmentally timed decay of an essential microRNA family is seed-sequence dependent. Cell Rep. 40, 111154 (2022).

39. Stubna, M. W., Shukla, A. & Bartel, D. P. Widespread destabilization of C. elegans microRNAs by the E3 ubiquitin ligase EBAX-1. RNA 31, rna.080276.124 (2024).

40. Seroussi, U. et al. A comprehensive survey of C. elegans argonaute proteins reveals organism-wide gene regulatory networks and functions. eLife 12, e83853 (2023).

41. Alberti, C. & Cochella, L. A framework for understanding the roles of miRNAs in animal development. Development 144, 2548–2559 (2017).

42. Dexheimer, P. J., Wang, J. & Cochella, L. Two MicroRNAs Are Sufficient for Embryonic Patterning in C. elegans. Curr. Biol. 30, 5058–5065.e5 (2020).

43. Grishok, A. et al. Genes and Mechanisms Related to RNA Interference Regulate Expression of the Small Temporal RNAs that Control C. elegans Developmental Timing. Cell 106, 23–34 (2001).

44. Hebbar, S. et al. Functional identification of microRNA-centered complexes in C. elegans. Sci. Rep. 12, 7133 (2022).

45. Zisoulis, D. G., Kai, Z. S., Chang, R. K. & Pasquinelli, A. E. Autoregulation of microRNA biogenesis by let-7 and Argonaute. Nature 486, 541–544 (2012).

46. Alvarez-Saavedra, E. & Horvitz, H. R. Many Families of C. elegans MicroRNAs Are Not Essential for Development or Viability. Curr. Biol. 20, 367–373 (2010).

47. Jan, C. H., Friedman, R. C., Ruby, J. G. & Bartel, D. P. Formation, regulation and evolution of Caenorhabditis elegans 3′UTRs. Nature 469, 97–101 (2011).

48. McGeary, S. E. et al. The biochemical basis of microRNA targeting efficacy. Science 366, (2019).

49. Yarychkivska, O. et al. Non-apoptotic death of the C. elegans linker cell is primed by MYRF-1 activation of pqn-41/polyQ. bioRxiv 2025.12.08.693091 (2025) doi:10.64898/2025.12.08.693091.

50. Kim, H., Lee, Y.-Y. & Kim, V. N. The biogenesis and regulation of animal microRNAs. Nat. Rev. Mol. Cell Biol. 26, 276–296 (2025).

51. Kingston, E. R., Blodgett, L. W. & Bartel, D. P. Endogenous transcripts direct microRNA degradation in Drosophila, and this targeted degradation is required for proper embryonic development. Mol. Cell 82, 3872–3884.e9 (2022).

52. Jones, B. T. et al. Target-directed microRNA degradation regulates developmental microRNA expression and embryonic growth in mammals. Genes Dev. 37, 661–674 (2023).

53. Shi, C. Y. et al. ZSWIM8 destabilizes many murine microRNAs and is required for proper embryonic growth and development. Genome Res. 33, 1482–1496 (2023).

54. Grimme, A. L. et al. A lncRNA drives developmentally-timed decay of all members of an essential microRNA family. bioRxiv 2025.07.30.667716 (2025) doi:10.1101/2025.07.30.667716.

55. Quiobe, S. P. et al. EBAX-1/ZSWIM8 destabilizes miRNAs, resulting in transgenerational inheritance of a predatory trait. Sci. Adv. 11, eadu0875 (2025).

56. Sherrard, R. et al. miRNAs cooperate in apoptosis regulation during C. elegans development. Genes Dev 31, 209–222 (2017).

57. Tran, A. T. et al. MiR-35 buffers apoptosis thresholds in the C. elegans germline by antagonizing both MAPK and core apoptosis pathways. Cell Death Differ. 26, 2637–2651 (2019).

58. Yarychkivska, O. et al. CED-3 caspase promotes dismantling but not onset of non-apoptotic linker cell death in C. elegans. bioRxiv 2025.10.15.682583 (2025) doi:10.1101/2025.10.15.682583.

59. Brenner, S. THE GENETICS OF CAENORHABDITIS ELEGANS. Genetics 77, 71–94 (1974).

60. Eroglu, M., Yu, B. & Derry, W. B. Efficient CRISPR/Cas9 mediated large insertions using long single-stranded oligonucleotide donors in C. elegans. FEBS J. 290, 4429–4439 (2023).

61. Gibson, D. G. et al. Enzymatic assembly of DNA molecules up to several hundred kilobases. Nat. Methods 6, 343–345 (2009).

62. Ent, F. van den & Löwe, J. RF cloning: A restriction-free method for inserting target genes into plasmids. J. Biochem. Biophys. Methods 67, 67–74 (2006).

63. Mello, C. C., Kramer, J. M., Stinchcomb, D. & Ambros, V. Efficient gene transfer in C.elegans: extrachromosomal maintenance and integration of transforming sequences. EMBO J. 10, 3959–3970 (1991).

64. Kutscher, L. M. & Shaham, S. Forward and reverse mutagenesis in C. elegans. WormBook 1–26 (2014) doi:10.1895/wormbook.1.167.1.

65. Kamath, R. S. & Ahringer, J. Genome-wide RNAi screening in Caenorhabditis elegans. Methods 30, 313–321 (2003).

66. Kamath, R. S. et al. Systematic functional analysis of the Caenorhabditis elegans genome using RNAi. Nature 421, 231–237 (2003).

67. Schindelin, J. et al. Fiji: an open-source platform for biological-image analysis. Nat. Methods 9, 676–682 (2012).

68. Lundquist, E. A., Reddien, P. W., Hartwieg, E., Horvitz, H. R. & Bargmann, C. I. Three C. elegans Rac proteins and several alternative Rac regulators control axon guidance, cell migration and apoptotic cell phagocytosis. Development 128, 4475–88 (2001).

69. Edelstein, A., Amodaj, N., Hoover, K., Vale, R. & Stuurman, N. Computer Control of Microscopes Using µManager. Curr. Protoc. Mol. Biol. 92, 14.20.1–14.20.17 (2010).

70. Meng, E. C. et al. UCSF ChimeraX: Tools for structure building and analysis. Protein Sci. 32, e4792 (2023).

