## Supplemental Figures and Tables for "*C. elegans* E3 ubiquitin ligase EBAX-1 promotes non-apoptotic linker cell-type death through target-directed miRNA degradation"

**SUPPLEMENTARY FIGURES AND LEGENDS**

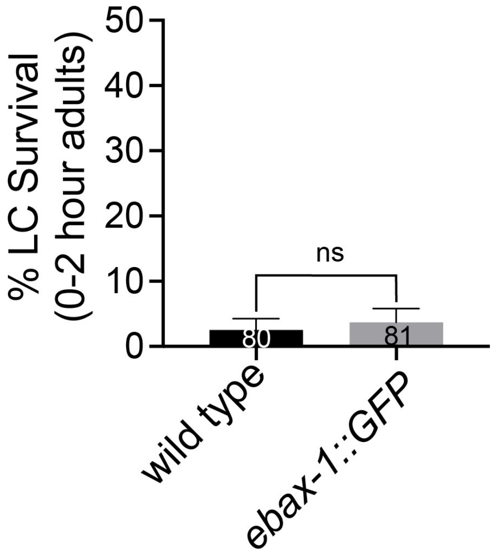


**Supplementary Figure 1. Insertion of *gfp* coding sequences at the C-terminal end of the endogenous *ebax-1* genomic locus does not affect linker cell death.**

Linker cell survival in indicated genotypes. ns, not significant; Fisher's exact test. Bar graph data are plotted as mean ± standard error of the proportion. Number of animals scored listed inside box.

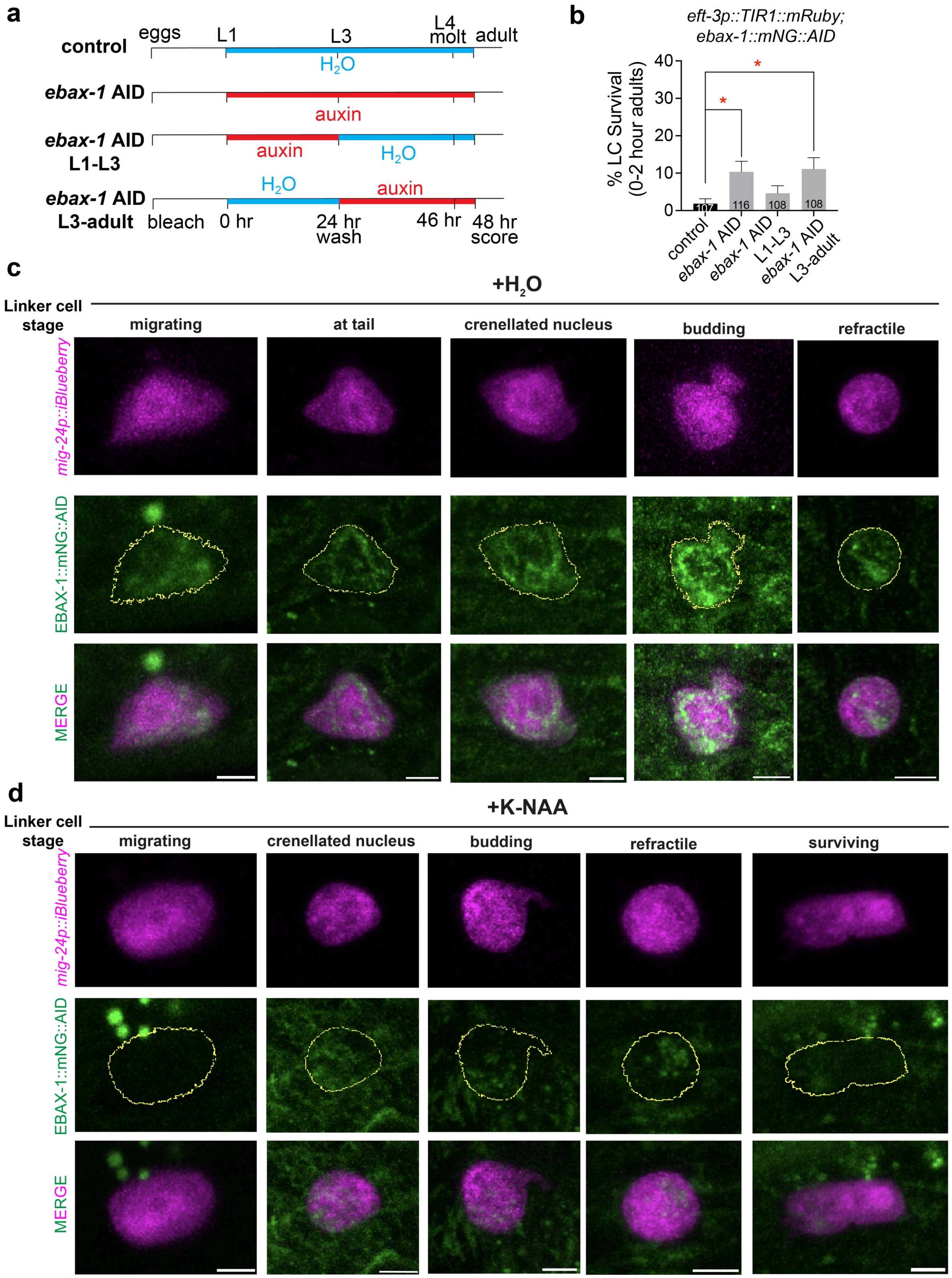


**Supplementary Figure 2. EBAX-1 likely promotes LCD during late larval development**

**(a)** Timeline of EBAX-1 auxin inducible degradation experiment, showing conditions relevant for **(b)**.

**(b)** Linker cell survival in *eft-3p::TIR1::mRuby; ebax-1::mNG::AID* mutant animals raised on water or auxin according to the conditions described in (A). (*) $p$ = 0.0113 for *ebax-1* AID, (*) $p$ = 0.0103 for *ebax-1* AID L3-adult*;* Fisher's exact test. Bar graph data are plotted as mean ± standard error of the proportion. Number of animals scored listed inside box.

**(c)** Representative confocal maximum intensity projections of linker cells at indicated stages in *eft-3p::TIR1::mRuby* animals expressing EBAX-1::mNeonGreen::AID and the linker cell marker *mig-24p::iBlueberry* raised on water. Images acquired using Airyscan mode. Maximum projections generated from 10 brightest *mig-24p::iBlueberry* slices from the z-stack, except for the budding condition which was a maximum projection of all slices in the z-stack to visualize the entire budding cell. Scale bar, 3 μm.

**(d)** Same as **(c)**, except raised on auxin. Scale bar, 3 μm.

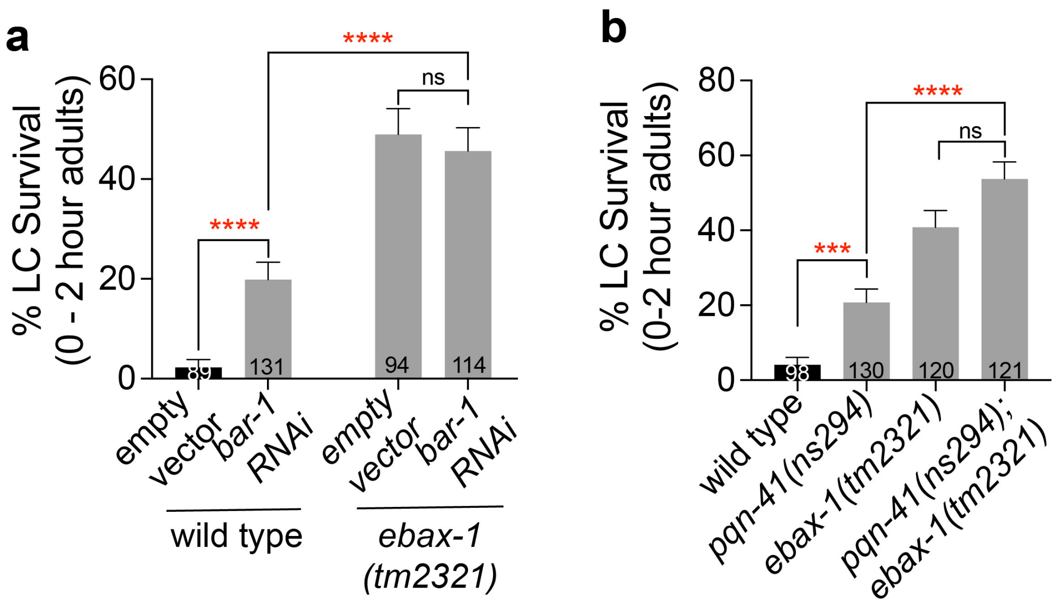


**Supplementary Figure 3. EBAX-1 functions downstream of the main activators of linker cell death**

**(a)** Linker cell survival in the wild type and in *ebax-1(tm2321)* mutant animals treated with empty vector and *bar-1 RNAi* constructs. (****) $p < 0.0001$, (ns) not significant $p = 0.6765$, Fisher's exact test.

**(b)** Linker cell survival in indicated genotypes. (***) $p = 0.0002$, (****) $p < 0.0001$, (ns) not significant $p = 0.0532$, Fisher's exact test.

Bar graph data are plotted as mean ± standard error of the proportion. Number of animals scored listed inside box. (ns) not significant.

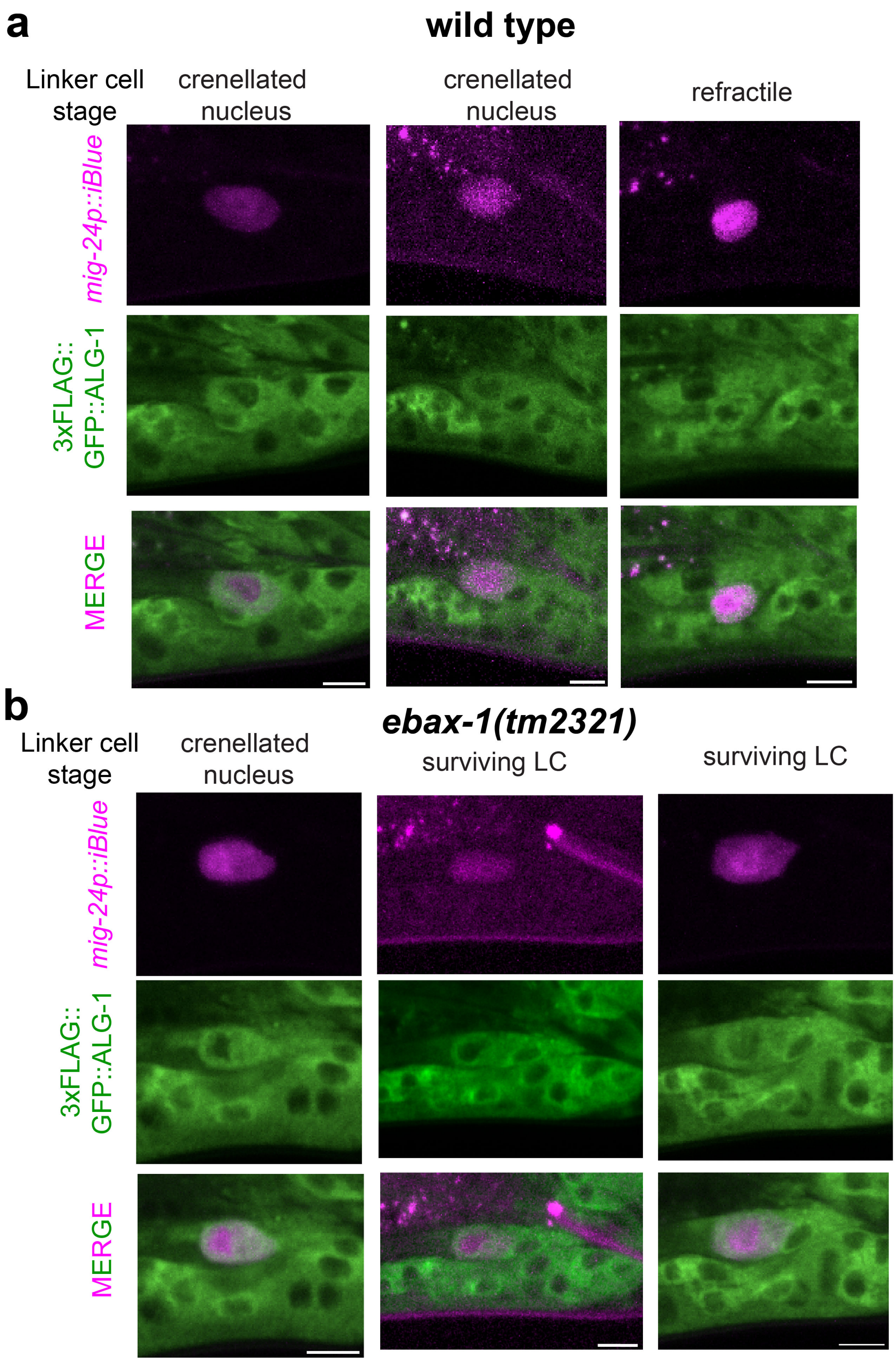


**Supplementary Figure 4. ALG-1 is expressed throughout the lifetime of the linker cell**

Representative confocal z-stack projections of wild type **(a)** and *ebax-1(tm2321)* **(b)** linker cells expressing 3xFLAG::GFP::ALG-1 and *mig-24p::iBlueberry* at indicated stages. Images of *mig-24p::iBlueberrry* are maximum intensity projections and images of 3xFLAG::GFP::ALG-1 are sum-intensity projections of z-stack images of the linker cell. Scale bar, 5 µm.

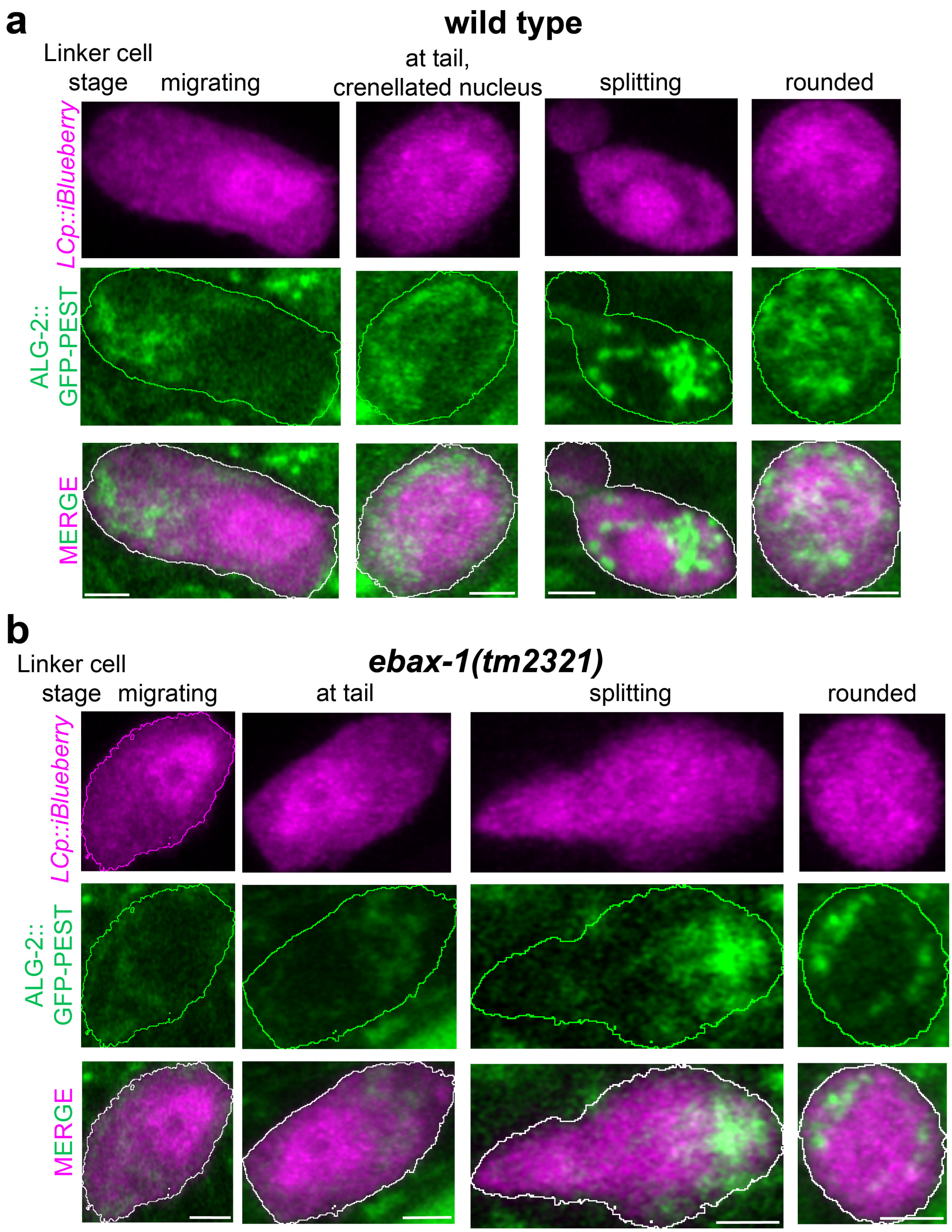


**Supplementary Figure 5. ALG-2 is expressed throughout the lifetime of the linker cell**

Representative confocal z-stack projections of wild type (**a**) and *ebax-1(tm2321)* mutant (**b**) linker cells at indicated stages expressing ALG-2::GFP-PEST and the linker cell marker *mig-24p::iBlueberry*. Images were acquired using Airyscan mode. Images of *mig-24p::iBlueberry* are maximum intensity projections and images of ALG-2::GFP-PEST are sum-intensity projections. Scale bar, 2 μm.

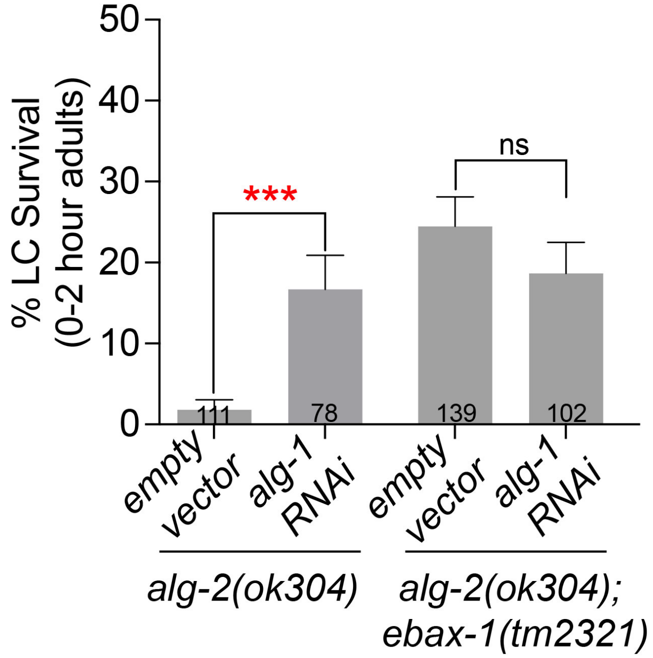


**Supplementary Figure 6. Loss of *alg-1* and *alg-2* does not have an additive effect on suppressing the linker cell survival defects of *ebax-1* mutants**

Linker cell survival in *alg-2(ok304)* and *alg-2(ok304); ebax-1(tm2321)* mutant animals treated with empty vector and *alg-1 RNAi*. (***) $p = 0.0002$, (ns) not significant $p = 0.3453$, Fisher's exact test. Bar graph data are plotted as mean ± standard error of the proportion. Number of animals scored listed inside box.

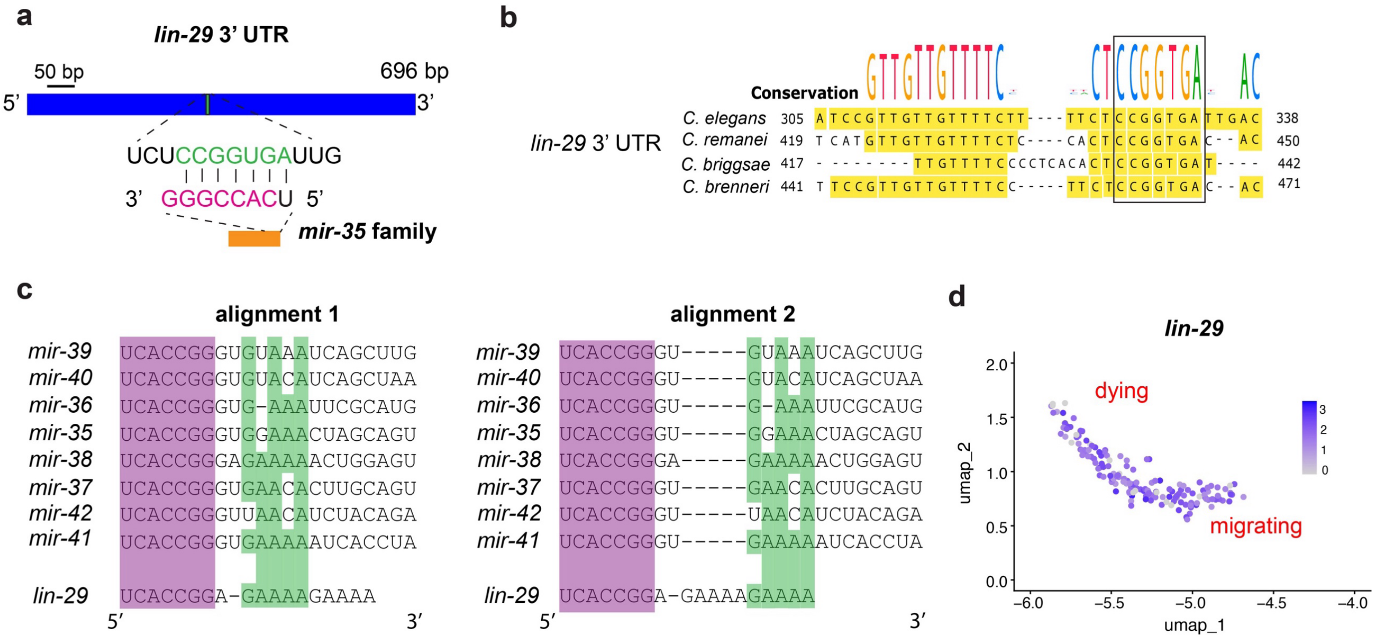


**Supplementary Figure 7. *lin-29* is a *mir-35* family target mRNA**

**(a)** Schematic of the 3' untranslated region (UTR) of *lin-29* showing the predicted sequence (green) recognized by the *mir-35* miRNA family seed sequence (pink).

**(b)** Sequence alignment of the *lin-29* 3' UTR containing the *mir-35* recognition site from indicated *Caenorhabditis* species. The conservation logo above the alignment represents nucleotide frequency at each position. Yellow shading highlights nucleotides that are identical to the *C. elegans lin-29* 3' UTR sequence. Boxed region marks the region complementary to the *mir-35* seed sequence.

**(c)** *mir-35* family miRNAs aligns with *lin-29* 3' UTR sequences outside the seed sequence in two alternative ways. Purple, seed sequence. Green, additional homology. The sequence shown for *lin-29* 3' UTR is the non-coding strand.

**(d)** *lin-29* expression in migrating and dying linker cells transcriptionally profiled using single-cell sequencing.

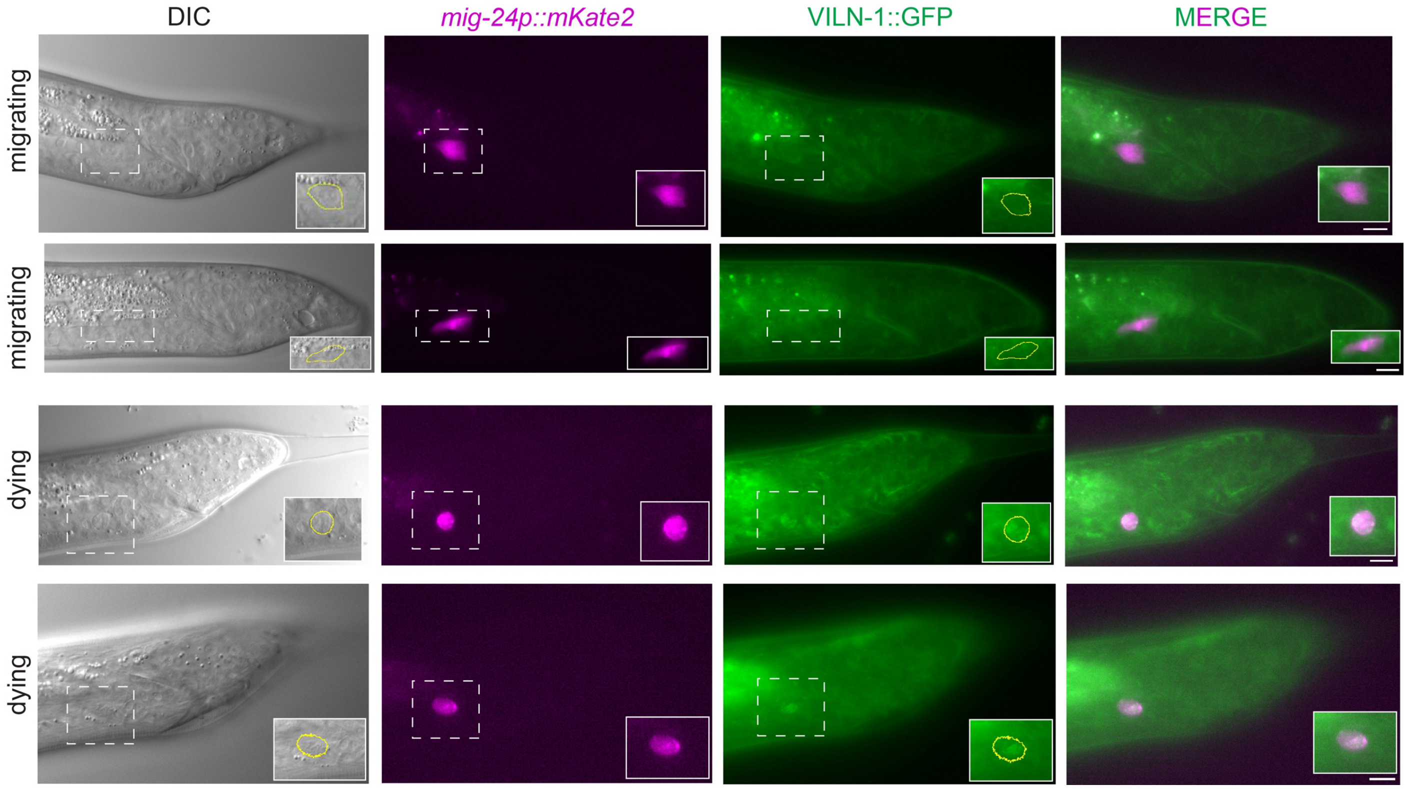


**Supplementary Figure 8. VILN-1 expression increases in dying linker cells.**

Representative DIC and fluorescent micrographs of a wild type animal expressing VILN-1::GFP during linker cell migration and death. The linker cell was visualized with *mig-24p::mKate2*. Inset, linker cell. Scale bar, 5 µm.

**SUPPLEMENTARY TABLES**

**Table S1: List of *C. elegans* strains used in this study**

| **Strain Name** | **Genotype** | **Source** |
|---|---|---|
| OS238 | *him-5(e1490) qIs56[lag-2p::GFP]* V | [1] |
| OS11663 | *ebax-1(tm2321)* IV*; him-5(e1490) qIs56[lag-2p::GFP]* V | This paper |
| OS11689 | *ebax-1(ju699)* IV*; him-5(e1490) qIs56[lag-2p::GFP]* V | This paper |
| OS11886 | *nsEx6035[* WRM062cC04(*ebax-1* fosmid, 5 ng/µL) + *odr-1p::RFP* (20 ng/µL) + *lag-2p::mCherry* (20 ng/µL) + *pBlueScript* (55 ng/µL)]; *ebax-1(tm2321)* IV*; him-5(e1490) qIs56[lag-2p::GFP]* V | This paper |
| OS13101 | *ebax-1(syb2784*[*ebax-1::GFP*]) IV; *him-5(e1490)* V; *nsIs650[mig-24p::mKate2* (10 ng/µL) + *unc-119*(+) (25 ng/µL) + *pBlueScript* (65 ng/µL)] X | This paper |
| OS13325 | *nsEx6475[mig-24p::ebax-1* (50 ng/µL) + *unc-122p::mCherry* (40 ng/µL)*]* line 1; *ebax-1(tm2321)* IV*; him-5(e1490) qIs56[lag-2p::GFP]* V | This paper |
| OS13326 | *nsEx6476[mig-24p::ebax-1* (50 ng/µL) + *unc-122p::mCherry* (40 ng/µL)*]* line 2; *ebax-1(tm2321)* IV*; him-5(e1490) qIs56[lag-2p::GFP]* V; | This paper |
| OS13327 | *nsEx6477[mig-24p::ebax-1* (50 ng/µL) + *unc-122p::mCherry* (40 ng/µL)*]* line 3; *ebax-1(tm2321)* IV*; him-5(e1490) qIs56[lag-2p::GFP]* V | This paper |
| OS13348 | *nsEx6484[lin-48p::ebax-1* (50 ng/µL) + *unc-122p::mCherry* (40 ng/µL)] line 1; *ebax-1(tm2321)* IV*; him-5(e1490) qIs56[lag-2p::GFP]* V | This paper |
| OS13349 | *nsEx6485[lin-48p::ebax-1* (50 ng/µL) + *unc-122p::mCherry* (40 ng/µL)] line 2; *ebax-1(tm2321)* IV*; him-5(e1490) qIs56[lag-2p::GFP]* V | This paper |
| OS13350 | *nsEx6486[lin-48p::ebax-1* (50 ng/µL) + *unc-122p::mCherry* (40 ng/µL)] line 3; *ebax-* | This paper |

| | | |
|---|---|---|
| | *1(tm2321)* IV*; him-5(e1490) qIs56[lag-2p::GFP]* V | |
| OS15318 | *ebax-1(syb7218[ebax-1::3xGAS::mNeonGreen::AID*]) IV; *him-5(e1490) qIs56[lag-2p::GFP] nsEx6485[lin-48p::ebax-1 + unc-122p::mCherry*] line 2; *ebax-1(tm2321)* IV*; him-5(e1490) qIs56[lag-2p::GFP]* V; *nsIs1045*[*mig-24p::TIR1::mRuby* (20 ng/µL) + *unc-122p::mCherry* (40 ng/µL)] X | This paper |
| OS13894 | *nsEx6788*[*mig-24p::ebax-1-N1(aa1-852)* (50 ng/µL) + *unc-122p::mCherry* (40 ng/µL)*]* line 1; *ebax-1(tm2321)* IV*; him-5(e1490) qIs56[lag-2p::GFP]* V | This paper |
| OS13895 | *nsEx6789*[*mig-24p::ebax-1-N1(aa1-852)* (50 ng/µL) + *unc-122p::mCherry* (40 ng/µL)] line 2; *ebax-1(tm2321)* IV*; him-5(e1490) qIs56[lag-2p::GFP]* V | This paper |
| OS13896 | *nsEx6790*[*mig-24p::ebax-1-N1(aa1-852)* (50 ng/µL) + *unc-122p::mCherry* (40 ng/µL)] line 3; *ebax-1(tm2321)* IV*; him-5(e1490) qIs56*[*lag-2p::GFP*] V | This paper |
| OS13898 | *nsEx6792*[*mig-24p::ebax-1-C1(aa853-1717)* (50 ng/µL) + *unc-122p::mCherry* (40 ng/µL)] line 1; *ebax-1(tm2321)* IV*; him-5(e1490) qIs56[lag-2p::GFP]* V | This paper |
| OS13900 | *nsEx6794*[*mig-24p::ebax-1-C1(aa853-1717)* (50 ng/µL) + *unc-122p::mCherry* (40 ng/µL)] line 2; *ebax-1(tm2321)* IV*; him-5(e1490) qIs56[lag-2p::GFP]* V | This paper |
| OS13899 | *nsEx6793*[*mig-24p::ebax-1-C1(aa853-1717)* (50 ng/µL) + *unc-122p::mCherry* (40 ng/µL)] line 3; *ebax-1(tm2321)* IV*; him-5(e1490) qIs56*[*lag-2p::GFP*] V | This paper |
| OS14329 | *nsEx7003[mig-24p::ebax-1-ΔBC-box + unc-122p::mCherry]* line 1; *ebax-1(tm2321)* IV*; him-5(e1490) qIs56[lag-2p::GFP]* V | This paper |
| OS14330 | *nsEx7004*[*mig-24p::ebax-1-ΔBC-box)* (50 ng/µL) + *unc-122p::mCherry* (40 ng/µL)] line 2; *ebax-1(tm2321)* IV*; him-5(e1490) qIs56*[*lag-2p::GFP*] V | This paper |
| OS14332 | *nsEx7006*[*mig-24p::ebax-1-ΔBC-box)* (50 ng/µL) + *unc-122p::mCherry* (40 ng/µL)] line 3; *ebax-1(tm2321)* IV*; him-5(e1490) qIs56[lag-2p::GFP]* V | This paper |

| | | |
|---|---|---|
| OS14474 | *nsEx7130*[*mig-24p::ebax-1-ΔCul2-box*(50 ng/µL) + *unc-122p::mCherry* (40 ng/µL)] line 1; *ebax-1(tm2321)* IV*; him-5(e1490) qIs56*[*lag-2p::GFP*] V | This paper |
| OS14475 | *nsEx7131*[*mig-24p::ebax-1-ΔCul2-box*(50 ng/µL) + *unc-122p::mCherry* (40 ng/µL)] line 2; *ebax-1(tm2321)* IV*; him-5(e1490) qIs56[lag-2p::GFP]* V | This paper |
| OS14476 | *nsEx7132*[*mig-24p::ebax-1-ΔCul2-box*(50 ng/µL) + *unc-122p::mCherry* (40 ng/µL)] line 3; *ebax-1(tm2321)* IV*; him-5(e1490) qIs56[lag-2p::GFP]* V | This paper |
| OS14333 | *nsEx7007*[*mig-24p::ebax-1-ΔSWIM* (50 ng/µL) + *unc-122p::mCherry* (40 ng/µL)] line 1; *ebax-1(tm2321)* IV*; him-5(e1490) qIs56[lag-2p::GFP]* V | This paper |
| OS14335 | *nsEx7009*[*mig-24p::ebax-1-ΔSWIM* (50 ng/µL) + *unc-122p::mCherry* (40 ng/µL)] line 2; *ebax-1(tm2321)* IV*; him-5(e1490) qIs56[lag-2p::GFP]* V | This paper |
| OS14334 | *nsEx7008*[*mig-24p::ebax-1-ΔSWIM*(50 ng/µL) + *unc-122p::mCherry* (40 ng/µL)] line 3; *ebax-1(tm2321)* IV*; him-5(e1490) qIs56[lag-2p::GFP]* V | This paper |
| OS14359 | *nsEx7029*[*mig-24p::ebax-1-ΔA*(50 ng/µL) + *unc-122p::mCherry* (40 ng/µL)] line 1; *ebax-1(tm2321)* IV*; him-5(e1490) qIs56[lag-2p::GFP]* V | This paper |
| OS14360 | *nsEx7030*[*mig-24p::ebax-1-ΔA*(50 ng/µL) + *unc-122p::mCherry* (40 ng/µL)] line 2; *ebax-1(tm2321)* IV*; him-5(e1490) qIs56[lag-2p::GFP]* V | This paper |
| OS14362 | *nsEx7032*[*mig-24p::ebax-1-ΔA*(50 ng/µL) + *unc-122p::mCherry* (40 ng/µL)] line 3; *ebax-1(tm2321)* IV*; him-5(e1490) qIs56[lag-2p::GFP]* V | This paper |
| OS15009 | *cul-2(or209)* III; *him-5(e1490) qIs56[lag-2p::GFP]* V | This paper |
| OS15010 | *cul-2(or209)* III; *ebax-1(tm2321)* IV; *him-5(e1490) qIs56[lag-2p::GFP]* V | This paper |
| OS7764 | *drSi28[hsf-1p::HSF-1(R145A)::GFP::unc-54 3' UTR + Cbr-unc-119(+)]* II; *him-5(e1490) qIs56[lag-2p::GFP]* V | [2] |
| OS4024 | *him-5(e1490) qIs56[lag-2p::GFP]* V; *sek-1(ag1)* X | [3] |

| | | |
|---|---|---|
| OS10135 | *drSi28[hsf-1p::HSF-1(R145A)::GFP::unc-54 3' UTR + Cbr-unc-119(+)]* II; *him-5(e1490) qIs56[lag-2p::GFP]* V; *sek-1(ag1)* X | [2] |
| OS13624 | *drSi28[hsf-1p::HSF-1(R145A)::GFP::unc-54 3' UTR + Cbr-unc-119(+)]* II; *ebax-1(tm2321)* IV; *him-5(e1490) qIs56[lag-2p::GFP]* V | This paper |
| OS9976 | *hsf-1(sy441) I*; *him-5(e1490) qIs56[lag-2p::GFP]* V | [2] |
| OS11949 | *hsf-1(sy441)* I; *ebax-1(ju699)* IV; *him-5(e1490) qIs56[lag-2p::GFP]* V | This paper |
| OS14614 | *tag-30(syb7669)* IV; *him-5(e1490) qIs56[lag-2p::GFP]* V | This paper |
| PHX7744 | *ebax-1(tm2321) tag-30(syb7669)* IV; *him-5(e1490) qIs56[lag-2p::GFP]* V | Suny Biotech |
| OS10580 | *let-70(ns770) IV; him-5(e1490) qIs56[lag-2p::GFP]* V | [2] |
| OS14038 | *ebax-1(ns1019)* IV; *him-5(e1490) qIs56[lag-2p::GFP]* V | This paper |
| OS13847 | *nT1[qIs51(IV;V)]/ebax-1(ns1019) let-70(ns770)* IV; *him-5(e1490) qIs56[lag-2p::GFP]* V | This paper |
| OS13143 | *ebax-1(syb2784*[*ebax-1::GFP*]) IV; *him-5(e1490)* V | This paper |
| OS4501 | *pqn-41(ns294)* III; *him-5(e1490) qIs56[lag-2p::GFP]* V | [3] |
| OS13787 | *pqn-41(ns294)* III; *ebax-1(tm2321)* IV; *him-5(e1490) qIs56[lag-2p::GFP]* V | This paper |
| OS14309 | *nsEx6991*[*mig-24p::ZSWIM8*(50 ng/µL) + *unc-122p::mCherry* (40 ng/µL)] line 1; *ebax-1(tm2321)* IV; *him-5(e1490) qIs56[lag-2p::GFP]* V | This paper |
| OS14310 | *nsEx6992*[*mig-24p::ZSWIM8*(50 ng/µL) + *unc-122p::mCherry* (40 ng/µL)] line 2; *ebax-1(tm2321)* IV; *him-5(e1490) qIs56[lag-2p::GFP]* V | This paper |
| OS14312 | *nsEx6994*[*mig-24p::ZSWIM8*(50 ng/µL) + *unc-122p::mCherry* (40 ng/µL)] line 3; *ebax-1(tm2321)* IV; *him-5(e1490) qIs56[lag-2p::GFP]* V | This paper |
| OS14041 | *alg-2(ok304)* II; *him-5(e1490) qIs56[lag-2p::GFP]* V | This paper |
| OS14039 | *alg-2(ok304)* II; *ebax-1(tm2321)* IV; *him-5(e1490) qIs56[lag-2p::GFP]* V | This paper |
| OS15348 | *drsh-1(lu82[myc::AID::3xFLAG::4xGGSG::drsh-1::4xGGSG::3xFLAG::AID::myc]*) I; *him-* | This paper |

| | | |
|---|---|---|
| | *5(e1490) qIs56[lag-2p::GFP]* V; *nsIs1045[mig-24p::TIR1::mRuby* (20 ng/µL) + *unc-122p::mCherry* (40 ng/µL)] X | |
| OS15350 | *drsh-1(luc82[myc::AID::3xFLAG::4xGGSG::drsh-1::4xGGSG::3xFLAG::AID::myc]*) I; *ebax-1(tm2321)* IV; *him-5(e1490) qIs56[lag-2p::GFP]* V; *nsIs1045[mig-24p::TIR1::mRuby*(20 ng/µL) + *unc-122p::mCherry* (40 ng/µL)*]* X | This paper |
| OS15431 | *pash-1(luc71[pash-1::2xGGSG::3xFLAG::AID::myc])* I; *him-5(e1490) qIs56[lag-2p::GFP]* V; *nsIs1045[mig-24p::TIR1::mRuby* (20 ng/µL) + *unc-122p::mCherry* (40 ng/µL)*]* X | This paper |
| OS15365 | *pash-1(luc71[pash-1::2xGGSG::3xFLAG::AID::myc])* I; *ebax-1(tm2321)* IV; *him-5(e1490) qIs56[lag-2p::GFP]* V; *nsIs1045[mig-24p::TIR1::mRuby* (20 ng/µL) + *unc-122p::mCherry* (40 ng/µL)*]* X | This paper |
| OS15265 | *nDf50[Δmir-35-41]* / *mln1[mIs14 dpy-10(e128)]* II; *him-5(e1490) qIs56[lag-2p::GFP]* V | This paper |
| OS15277 | *nDf50[Δmir-35-41]* / *mln1[mIs14 dpy-10(e128)]* II; *ebax-1(tm2321)* IV; *him-5(e1490) qIs56[lag-2p::GFP]* V | This paper |
| OS15163 | *nDf50[Δmir-35-41] nDf49[Δmir-42] / mln1[mIs14 dpy-10(e128)]* II; *him-5(e1490) qIs56[lag-2p::GFP]* V | This paper |
| OS15177 | *nDf50[Δmir-35-41] nDf49[Δmir-42] / mln1[mIs14 dpy-10(e128)]* II; *ebax-1(tm2321)* IV; *him-5(e1490) qIs56[lag-2p::GFP]* V | This paper |
| OS15433 | *nEx479*[*mig-24p::mir-35-41* (50 ng/uL) + *unc-122p::mCherry* (40 ng/uL)] line 1 *him-5(e1490) qIs56[lag-2p::GFP]* V | This paper |
| OS15434 | *nEx480*[*mig-24p::mir-35-41* (50 ng/uL) + *unc-122p::mCherry* (40 ng/uL)] line 2 *him-5(e1490) qIs56[lag-2p::GFP]* V | This paper |
| OS15491 | *viln-1(ns1112[viln-1::GFP])* I; *him-5(e1490)* V; *nsIs650*[*mig-24p:: mKate2* (10 ng/µL) + *unc-119*(+) (25 ng/µL) + *pBlueScript* (65 ng/µL)] X | This paper |
| OS7637 | *him-8(e1489) IV; rde-1(ne219) qIs56[lag-2p::GFP] V; nsIs387[mig-24p::rde-1::SLC-mCherry + lag-2p::mCherry]* | [2] |

| | | |
|---|---|---|
| OS15327 | *viln-1(ok2413)* I*; him-5(e1490) qIs56[lag-2p::GFP]* V | This paper |
| PHX8021 | *nsIs712[mig-24p::iBlueberry-P2A-HO1* (25 ng/µL) + *unc-119(+)* (25 ng/µL) + pBlueScript (50 ng/µL)*]* I; *alg-2(syb8021[alg-2::GFP-PEST])* II; *unc-119(ed3)* III; *him-5(e1490)* V | Suny Biotech |
| OS14962 | *nsIs712[mig-24p::iBlueberry-P2A-HO1* (25 ng/µL) + *unc-119(+)* (25 ng/µL) + pBlueScript (50 ng/µL)*]* I; *alg-2(syb8021[alg-2::GFP-PEST])* II; *unc-119(ed3)* III; *ebax-1(tm2321) IV*; *him-5(e1490)* V | This paper |
| OS14337 | *nsIs712[mig-24p::iBlueberry-P2A-HO1*(25 ng/µL) + *unc-119(+)* (25 ng/µL) + pBlueScript (50 ng/µL)*]* I; *him-5(e1490)* V; *alg-1(ap423[3xFLAG::GFP::ALG-1])* X | This paper |
| OS14466 | *nsIs712[mig-24p::iBlueberry-P2A-*(25 ng/µL) + *unc-119(+)* (25 ng/µL) + pBlueScript (50 ng/µL)*]* I; *ebax-1(tm2321)* IV; *him-5(e1490)* V; *alg-1(ap423[3xFLAG::GFP::ALG-1])* X | This paper |

**Table S2: Oligonucleotides used in this study**

| **Name** | **Description** | **Sequence** |
|---|---|---|
| *ebax-1(ns1019)* crRNA 1 | Left crRNA to generate *ebax-1(ns1019)* deletion | AACACTTTTCTCTGTGAGG |
| *ebax-1(ns1019)* crRNA 2 | Right crRNA to generate *ebax-1(ns1019)* deletion | GTTGGAGGAACACTTTGAGC |
| *ebax-1(ns1019)* ssODN | ssODN template to generate a 909 bp deletion in the fifth exon of *ebax-1* | CAAATCTATCGAACAGTTACAAGATGGTTCCACTTCAAAGTGTTCCTCCAACTGCTTCACCTGCTCTTCA |
| *viln-1* crRNA | crRNA that cuts at the end of the *viln-1* coding sequence | AACGATGCACGGAAAAGAGT |
| LBH 212 | F primer to amplify linker-GFP with 35 bp overhangs homologous to *viln-1* coding sequence immediately before the STOP codon. Homology region contains 4 silent mutations to prevent recutting by the crRNA: ACGGAAAAGA is changed to CCGGAAGCGC | AACAGAACGATGCCCGGAAGCGCGTTGGACTATTCGGAGCATCGGGAGCC |
| LBH 213 | R primer to amplify linker-GFP with 35 bp overhangs homologous to the *viln-1* 3' UTR immediately after the STOP codon. | cgcctctagtgagtcaatataaataaaaaatatgtCTATTTGTATAGTTCATCCATGCC |
| LBH 5 | F primer to amplify *ebax-1*cDNA fragment 1 with overhangs to place into pPD49.26 fire vector | aacattttcaggaggacccttggctagcATGCTGCCAAATGCAGTGGG |
| LBH 6 | R primer to amplify *ebax-1* cDNA fragment 1 | ACGCACGTTCTCGAATTTTTCG |
| LBH 7 | F primer to amplify *ebax-1* cDNA fragment 2 | TGTCCCGATGAGTGATACCC |
| LBH 8 | R primer to amplify *ebax-1* cDNA fragment 2 with overhangs to place into pPD49.26 fire vector | ctcagatatcaataccatggtaccgtcgacCTATAATGGGCTGAGTTGCCC |
| NR 700 | F primer to amplify pPD49.26 fire vector | gtcgacggtaccatggtattgatatc |
| NR 701 | R primer to amplify pPD49.26 fire vector | gctagccaagggtcctcctg |
| LBH 17 | F primer to amplify *ebax-1* cDNA with homology arms to *mig-24* promoter vector backbone | GTCATCTTACACAATTTTATTAAACCCGGGATGCTGCCAAATGCAGTGGG |

| | | |
|---|---|---|
| LBH 18 | R primer to amplify *ebax-1* cDNA with homology arms to *mig-24* and *lin-48* vector backbone (3' to insertion) | GTAGCGACCGGCGCTCAGTTGGAATTCttaCTATAATGGGCTGAGTTGCCC |
| LBH 15 | F primer to amplify *mig-24* and *lin-48* promoter vector backbone | taaGAATTCCAACTGAGCGCCG |
| LBH 16 | R primer to amplify *mig-24* promoter vector backbone | CCCGGGTTTAATAAAATTGTGTAAGA |
| LBH 19 | R primer to amplify *lin-48* promoter vector backbone (use with LBH 14) | cccgggctgaaattgagcaga |
| LBH 57 | R primer to amplify *ebax-1*-N1(aa1-852) with overhangs to put into *mig-24* promoter (use with LBH 17) | GTAGCGACCGGCGCTCAGTTGGAATTCttaCTACAAAATGCCAACAAATTCCGTCT |
| LBH 58 | F primer to amplify *ebax-1*-C1(aa853-1717) with overhangs to put into *mig-24* promoter (LBH 18) | GTCATCTTACACAATTTTATTAAACCCGGGATGGATCAAATCTGGCAAACTGT |
| LBH 67 | F primer to amplify *mig-24p::ebax-1*-ΔSWIM | TATGATGATAATTCATCTTTCCATTACGAAAGGAGCAATATAAAATTC |
| LBH 68 | R primer to amplify *mig-24p::ebax-1*-ΔSWIM | GAATTTTATATTGCTCCTTTCGTAATGGAAAGATGAATTATCATCATA |
| LBH 63 | F primer to amplify *mig-24p::ebax-1*-ΔBC-box | AGAGGAGATGATTCAATTGGAACTTCGTTTCAGCAACTTGAAGAAATG |
| LBH 64 | R primer to amplify *mig-24p::ebax-1*-ΔBC-box | CATTTCTTCAAGTTGCTGAAACGAAGTTCCAATTGAATCATCTCCTCT |
| LBH 69 | F primer to amplify *mig-24p::ebax-1*-ΔA | ATCAATTCAATGAGTGATTTTGATCTGTCATCTCAGAATATGAAGTTC |
| LBH 70 | R primer to amplify *mig-24p::ebax-1*-ΔA | GAACTTCATATTCTGAGATGACAGATCAAAATCACTCATTGAATTGAT |
| LBH 65 | F primer to amplify *mig-24p::ebax-1*-ΔCul2-box with Q5 mutagenesis site-directed kit | ATAGTTCATTGTTTCCCACAATC |
| LBH 66 | R primer to amplify *mig-24p::ebax-1*-ΔCul2-box with Q5 mutagenesis site-directed kit | AGTGGAGATTGGTGGAGG |
| LBH 86 | F primer to amplify *zswim8* cDNA with overhangs to put into *mig-24* promoter | GTCATCTTACACAATTTTATTAAACCCGGGATGGAGCTGATGTTTGCAGA |

| | | |
|---|---|---|
| LBH 87 | R primer to amplify *zswim8* cDNA with overhangs to put into *mig-24* promoter | GTAGCGACCGGCGCTCAGTTGGAATTCttaTCAGGGGGAGAAGGTGGC |
| LBH 124 | F primer to amplify *TIR1::mRuby* with overhangs to put into *mig-24* promoter | GTCATCTTACACAATTTTATTAAACCCGGGATGCAAAAGAGAATCGCCTTG |
| LBH 125 | R primer to amplify *TIR1::mRuby* with overhangs to put into *mig-24* promoter | GTAGCGACCGGCGCTCAGTTGGAATTCttaTTATCCTCCTCCAAGTCCAGC |
| LBH 201 | F primer to amplify *mir-35-41* genomic sequence with overhangs to put into *mig-24* promoter | GTCATCTTACACAATTTTATTAAACCCGGGgtgtcaaatgattacgacga |
| LBH 202 | R primer to amplify *mir-35-41* genomic sequence with overhangs to put into *mig-24* promoter | GTAGCGACCGGCGCTCAGTTGGAATTCttacggaaaggtacatatgcgcc |

**Table S3: Plasmids Used in this Study**

| **Plasmid Name** | **Description** | **Notes** |
|---|---|---|
| pLBH63 | *ebax-1* cDNA in pPD49.26 Fire expression vector | *ebax-1* cDNA was amplified from *C. elegans* cDNA as two overlapping fragments (fragment 1: aa 1- 872, and fragment 2: aa 860 – 1717) with primers (fragment 1 - LBH5/6, fragment 2 – LBH 7/8) that contained homology arms for insertion into the pPD49.26. pPD49.26 was amplified using NR700 and NR701. pPD49.26 was purchased from Addgene. |
| pLBH65 | *mig-24p::ebax-1* cDNA | *ebax-1* cDNA was amplified from pLBH63 using primers with overhangs homologous to a plasmid containing the *mig-24* promoter (LBH17/18). The *mig-24* promoter vector backbone was amplified from pLMK113 (*mig24p::mKate2*) using primers LBH15/16. |
| pLBH66 | *lin-48p::ebax-1* cDNA | *ebax-1* cDNA was amplified from pLBH63 using primers with overhangs homologous to the vector containing the *lin-48* promoter (LBH20/18). The *lin-48* promoter vector backbone was amplified from pLMK21 (*lin-48p::mKate2::rab-35*) using primers LBH15/19 |
| pLBH67 | *mig-24p::ebax-1-N1*(aa1-852) | *ebax-1-N1*(aa 1- 852) was amplified from pLBH63 with primers homologous to the *mig-24* promoter vector backbone (LBH 17/57). The vector containing the *mig-24* promoter was amplified as described for pLBH65. |
| pLBH69 | *mig-24p::ebax-1-C1*(aa853-1717) | *ebax-1-C1*(aa853-1717) was amplified from pLBH63 with primers homologous to the |

| | | |
|---|---|---|
| | | *mig-24* promoter vector backbone (LBH 58/18). The vector containing the *mig-24* promoter was amplified as described for pLBH65. |
| pLBH70 | *mig-24p::ebax-1-ΔSWIM* | Cloned from pLBH65 (*mig-24p::ebax-1*) by amplifying entire plasmid except for the predicted SWIM domain in *ebax-1* cDNA using primers LBH67/68. |
| pLBH71 | *mig-24p::ebax-1-ΔBC-box* | Cloned from pLBH65( *mig-24p::ebax-1*) by amplifying entire plasmid except for the predicted BC-box in *ebax-1* cDNA using primers LBH 63/64. |
| pLBH72 | *mig-24p::ebax-1-ΔA* | Cloned from pLBH65( *mig-24p::ebax-1*) by amplifying entire plasmid except for the predicted A domain in *ebax-1* cDNA using primers LBH69/70. |
| pLBH73 | *mig-24p::zswim8* | *zswim8* cDNA was amplified from pCMV-sport6-zswim8 (transOMIC technologies) using primers homologous to the *mig-24* vector backbone (LBH86/87). The vector containing the *mig-24* promoter was amplified as described for pLBH65. |
| pLBH75 | *mig-24p::ebax-1-ΔCul2-box* | Cloned from pLBH65 (*mig-24p::ebax-1*) using primers LBH 65/66 and the Q5 site-directed mutagenesis kit (NEB). |
| pLBH78 | *mig-24p::TIR1::mRuby* | *TIR1::mRuby* was amplified from pLZ31(*eft-3p::TIR1::mRuby*) using primers homologous to the *mig-24* vector backbone (LBH124/125). The vector containing the *mig-24* promoter was amplified as described for pLBH65. |

| | | |
|---|---|---|
| pLBH86 | *mig-24p::mir-35-41* | *mir-35-41* was amplified from pKM13(pCFJ352-*mir-35-41* rescue donor) using primers homologous to the *mig-24* vector backbone (LBH201/202). The vector containing the *mig-24* promoter was amplified as described for pLBH65. |

### SUPPLEMENTARY REFERENCES


1. Abraham, M. C., Lu, Y. & Shaham, S. A morphologically conserved nonapoptotic program promotes linker cell death in Caenorhabditis elegans. *Dev Cell* **12**, 73–86 (2007).

2. Kinet, M. J. *et al.* HSF-1 activates the ubiquitin proteasome system to promote non-apoptotic developmental cell death in C. elegans. *Elife* **5**, (2016).

3. Blum, E. S., Abraham, M. C., Yoshimura, S., Lu, Y. & Shaham, S. Control of nonapoptotic developmental cell death in Caenorhabditis elegans by a polyglutamine-repeat protein. *Science* **335**, 970–3 (2012).
